# Multi-scale systems genomics analysis predicts pathways, cell types and drug targets involved in normative human cognition variation

**DOI:** 10.1101/751933

**Authors:** Shraddha Pai, Shirley Hui, Philipp Weber, Soumil Narayan, Owen Whitley, Peipei Li, Viviane Labrie, Jan Baumbach, Gary D Bader

## Abstract

An open challenge in human genetics is to better understand the link between genotype variation and the various molecular, cellular, anatomical and physiological systems that it can affect. To address this challenge, we performed genotype-phenotype-systems analysis for accuracy in nine cognitive tasks from the Philadelphia Neurodevelopmental Cohort (3,319 individuals aged 8-21 years). We report a region of genome-wide significance within the 3’ end of the *FBLN1* gene (p=4.6×10^−8^), associated with nonverbal reasoning, a heritable form of complex reasoning ability. Integration of published brain-specific ‘omic maps reveals that *FBLN1* shows greatest expression in the fetal brain, is a marker of neural progenitor cells, is differentially expressed in schizophrenia and increases genetic risk for bipolar disorder. These findings suggest that nonverbal reasoning and *FBLN1* variation warrant further investigation in studies of neurodevelopmental disorders and psychosis. Using genotype-pathway analysis, we identify pathways related to development and to autonomic nervous system dysfunction associated with working memory accuracy. Top-ranking pathway genes include those genetically associated with multiple diseases with working memory deficits, such as schizophrenia and Parkinson’s disease, and that are also markers for specific brain cell types. Our findings identify novel molecular players involved in specific cognitive tasks and link variants to genes, pathways, cell types, diseases and drugs. This work advances the “molecules-to-behaviour” view of cognition, and provides a framework for using systems-level organization of data for other biomedical domains.

## Introduction

The growth in genomics and functional annotation resources over the past decade provides an opportunity to build models of how changing genotype affects multiple levels of system organization underlying a phenotype, from genes and molecules through to pathway, cell, cell circuit, anatomy and physiology system levels (systems-genomics analysis). This opportunity complements a conceptual shift to systems-level thinking in many biomedical fields. For example, a major drive in psychiatry is the reconceptualization of mental illnesses as brain disorders treatable by neurobiological system-grounded therapies, such as working memory deficits in schizophrenia^1^. As a shared guide for the field, the U.S. National Institute of Mental Health has developed a “genes-to-behaviour” framework that deconstructs human behaviour into neurobehavioural domains, such as cognition and social processing^2^. Each of these constructs has subconstructs and these are linked to a variety of systems level concepts. While the genetic architecture of overall cognitive ability (i.e., intelligence) has been studied by large-scale GWAS^3-5^, little is known about the molecular basis of more detailed neurocognitive phenotypes.

In this work, we identify genetic variants associated with normative variation in nine cognitive phenotypes measured in the Philadelphia Neurodevelopmental Cohort (PNC). We selected this study for our systems-genomics analysis as the phenotypes measured were designed around systems-level neuroscience theory. For example, tasks requiring use of working memory, a type of short-term memory that recruits a cortical-subcortical network including the dorsolateral prefrontal cortex, shows a genetic component in twins, and is impaired in schizophrenia ^6-8^. Thus, we hypothesize that this data will yield systems-genomics signal, that is genetic variants linked to one or more system level scales of phenotype-related organization. To our knowledge, there have been no reports of genotype-phenotype analyses on the PNC dataset. Using diverse functional genomics resources, we link variants to genes, pathways, brain cell types, brain systems, predicted drug targets, and diseases, providing a systems-level view of the genetics of the neurodevelopmental phenotypes under study. With standardized and well-controlled cognitive tests and genotyping on over 8,000 community youths aged 8-21 years, the PNC is the largest publicly-available dataset for genotype-phenotype analysis of developmental cognition^9,10^. All participants have computerized neurocognitive test battery scores, measuring speed and accuracy in multiple cognitive domains. These measures have neurobehavioural validity^11^, SNP-based heritability^12^, and disease relevance^11,13^. Multiple cognitive test scores in the PNC demonstrate significant SNP-based heritability^12^, and reduced test scores are correlated with increased genetic risk of psychiatric disease^13^. Moreover, the PNC captures the age range through which some cognitive abilities, such as working memory, mature to stable adult levels^14,15^. Despite the relatively small size of this dataset by GWAS standards, we reasoned that the PNC dataset provides a valuable opportunity to study the molecular and systems basis of cognitive tasks impaired in disease and evaluate how a systems-genomics approach can increase statistical and interpretive power compared to standard SNP and gene-based analysis approaches, both of which are performed here to enable us to compare these approaches.

## Methods

Cognitive assessment was performed using the Penn Computerized Neurocognitive Battery (CNB), which was customized and shortened for a pediatric population. Performance is measured by a session of trials containing items with varying levels of difficulty, which allows the test to capture nuances in speed and accuracy measures. Tests were also developed through evaluation by psychological investigators to ensure tasks could measure the phenotype of interest (i.e. had “construct validity”) and reliability between test-takers and through retakes^11^.

### Genetic imputation

The samples (n=8,719) were all genotyped using Illumina or Affymetrix SNP-array platforms by the Center for Applied Genomics at The Children’s Hospital of Philadelphia.^16^ The workflow for genomic imputation is shown in Supplementary Figure 1. Genotypes for the four most frequent microarray genotyping platforms were downloaded from dbGaP (phs000607.v1). We performed genetic imputation for the Illumina Human610-Quad BeadChip, the Illumina HumanHap550 Genotyping BeadChip v1.1, Illumina HumanHap550 Genotyping BeadChip v3, and the Affymetrix AxiomExpress platform (Supplementary Table 1, total of 6,502 samples before imputation), using the protocol recommended by the EMERGE consortium^17^. Imputation was performed as follows:

#### Step 1: Platform-specific plink quality control

Quality control was first performed for each microarray platform separately. Single nucleotide polymorphisms (SNPs) were limited to those on chr1-22. SNPs in linkage disequilibrium (LD) were excluded (--indep-pairwise 50 5 0.2), and alleles were recoded from numeric to letter (ACGT) coding. Samples were excluded if they demonstrated heterozygosity > 3 standard deviations (SD) from the mean, or if they were missing >=5% genotypes. Where samples had pairwise Identity by Descent (IBD) > 0.185, one of the pair was excluded. Variants with minor allele frequency (MAF) < 0.05 were excluded, as were those failing Hardy-Weinberg equilibrium with p < 1e-6 and those missing in >=5% samples.

#### Step 2: Convert coordinates to hg19

LiftOver^18^ was used to convert SNPs from human genome assembly version hg18 to hg19; Hap550K v1 data was in hg17 and was converted from this build to hg19.

#### Step 3: Strand-match check and prephasing

ShapeIt v2.r790^19^ was used to confirm that the allelic strand in the input data matched that in the reference panel; where it did not, allele strands were flipped (shapeit “–check” flag). ShapeIt was used to prephase the variants using the genetic_b37 reference panel (downloaded from the Shapeit website, http://www.shapeit.fr/files/genetic_map_b37.tar.gz)

#### Step 4: Imputation

Genotypes were imputed using Impute2 v2.3.2^20^ and a reference panel from the 1,000 Genomes (phase 1, prephased with Shapeit2, no singletons, 16 June 2014 release, downloaded from https://mathgen.stats.ox.ac.uk/impute/data_download_1000G_phase1_integrated_SHAPEIT2_16-06-14.html) was used for imputation, using the parameter settings “–use_prephased_g –Ne 20000 –seed 367946”. Average concordance for all chromosomes was ∼95%, indicating successful imputation (Supplementary Figure 2). Imputed genotypes were merged across all platforms using software from the Ritchie lab^17^ (impute2-group-join.py, from https://ritchielab.org/software/imputation-download) and converted to plink format. Following previous PNC genotype analysis^12^, only SNPs with info score > 0.6 were retained, and deletions/insertions were excluded (plink “-snps-only just-acgt” flags). As preliminary quality control, when merging across chromosomes, samples with missingness exceeding 99% were excluded, as were SNPs with MAF < 1% and with missingness exceeding 99%. This step resulted in 10,845,339 SNPs and 6,327 individuals.

#### Step 5: Post-imputation quality control

The HapMap3 panel was used to assign genetic ancestry for samples, using steps from ^21^ (Supplementary Figure 3). Individuals within 5 SD of the centroid of the HapMap3 CEU (Utah residents with Northern or Western European ancestry) or TSI (Tuscans in Italy) clusters were assigned to belong to the respective groups, and were classified as being of European descent; 3,441 individuals pass this filter. Individuals with >5% missing data were excluded, as was one of each pair of individuals with IBS > 0.185 (47 individuals); 3,394 individuals passed this filter. Variants that were symmetric or in regions of high LD (Supplementary Table 2) were excluded (9,631,316 SNPs passed). Variants with >5% missingness were excluded (1,569,407 SNPs excluded). Finally, SNPs with MAF < 0.01 (3,168,339 SNPs) and failing Hardy-Weinberg equilibrium (HWE) with p value < 1e-6 (373 SNPs) were excluded, resulting in 4,893,197 SNPs. As only high-quality SNPs were retained after imputation, post-processing steps were performed only once. In sum, the imputation process resulted in 3,394 individuals and 4,893,197 SNPs available for downstream analysis.

### Phenotype processing

Phenotype data was downloaded from dbGaP for 8,719 individuals. 637 individuals with severe medical conditions (Medical rating=4) were excluded to avoid confounding the symptoms of their conditions with performance on the cognitive tests^12^. Linear regression was used to regress out the effect of age at test time (variable name: “age at cnb”) and sex from sample-level phenotype scores, and the residualized phenotype was used for downstream analysis.

The nine phenotypes selected for systems-genomics analysis are measures of overall performance accuracy in the Penn Computerized Neurocognitive Test Battery (CNB; Supplementary Table 3) and represent major cognitive domains. Tasks mapped to domains in the following manner: verbal reasoning, nonverbal reasoning, and spatial reasoning measured complex cognition; attention allocation and working memory measured executive function; recall tests for faces, words and objects measured declarative memory, and emotion identification measured social processing. Following regression, none of the phenotypes were significantly correlated with age after Bonferroni correction, indicating that the age effect had been reduced (Supplementary Table 4). Following guidelines from previous analyses on these data^13^, individuals with scores more than four standard deviations from the mean for a particular test, representing outliers, were excluded from the analysis of the corresponding phenotype. For a given phenotype, only samples with a code indicating a valid test score (codes “V” or “V2”) were included; e.g. for pfmt_tp (Penn Face Memory Test), only samples with pfmt_valid = “V” or “V2” were retained; the rest had scores set to NA. Finally, each phenotype was dichotomized so that samples in the bottom 33^rd^ percentile were relabeled as “poor” performers and those in the top 33^rd^ were set to be “good” performers; for a given phenotype, this process resulted in ∼1,000 samples in each group (Supplementary Table 3). Where an individual had good or poor performance in multiple phenotypes, they were included in the corresponding group for each of those phenotypes.

### Genetic association analysis

For each of nine CNB phenotypes, marginal SNP-level association was calculated using a mixed-effects linear model (MLMA), using the leave-one-chromosome-out (LOCO) method of estimating polygenic contribution (GCTA v1.97.7beta software^22^). In this strategy, a mixed-effect model is fit for each SNP:

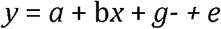

where *y* is the binarized label (good/poor performer on a particular task), *x* measures the effect of genotype (indicator variable coded as 0, 1 or 2), *g*-represents the polygenic contribution of all the SNPs in the genome (here, the ∼4.89M imputed SNPs), and *e* represents a vector of residual effects. In the LOCO variation, *g*-is calculated using a chromosome-specific genetic relatedness matrix, one that excludes the chromosome on which the candidate SNP is located^22^. SNPs and associated genes were annotated as described in Supplementary Notes 1-4.

### Hi-C Data Processing

We downloaded publicly-available Hi-C data from human prefrontal cortex tissue^23,24^ (Illumina HiSeq 2000 paired-end raw sequence reads; n=1 sample; 746 Million reads; accession: GSM2322542 [https://www.ncbi.nlm.nih.gov/geo/query/acc.cgi?acc=GSM2322542]). We used Trim Galore^25^ (v0.4.3) for adapter trimming, HICUP^26^ (v0.5.9) for mapping and performing quality control, and GOTHIC^27^ for identifying significant interactions (Bonferroni p <0.05), with a 40 kb resolution. Hi-C gene annotation involved identifying interactions with gene promoters, defined as ± 2 kb of a gene TSS. This analysis identified 303,464 DNA-DNA interactions used for our study.

### SNP to gene mapping for annotation and enrichment analyses

SNPs were mapped to genes using a combination of genome position information (i.e. closest gene), brain-specific expression Quantitative Trait Locus (eQTL) and higher-order chromatin interaction (Hi-C) information. Gene definitions were downloaded from Gencode (ftp://ftp.ebi.ac.uk/pub/databases/gencode/Gencode_human/release_32/GRCh37_mapping/gencode.v32lift37.basic.annotation.gtf.gz). Only genes with “protein_coding” biotype were included (20,076 unique gene symbols), to simplify interpretation of cellular mechanisms using pathway annotation information, which almost completely include only protein coding genes. Using chromatin state maps from the Roadmap Epigenomics project^28^, we compiled a list of open chromatin and enhancer regions in brain tissue. These comprised maps derived from 13 human brain samples, including: neurospheres, angular gyrus, anterior caudate, germinal matrix, hippocampus, inferior temporal lobe, dorsolateral prefrontal cortex, substantia nigra, and fetal brain of both sexes (samples E053, E054, E067, E068, E069, E070, E071, E072, E073, E074, E081, E082, and E125), downloaded from http://www.roadmapepigenomics.org/. Open chromatin states were defined as genomic regions with epigenomic roadmap project’s core 15-state model values <=7. Enhancers were defined as those labeled with states “Enh” and “EnhG”.

For eQTL-based mapping, we searched for significant eQTLs in 12 types of brain tissue (GTEx v7: Amygdala, Anterior cingulate cortex BA24, Caudate basal ganglia, Cerebellar Hemisphere, Cerebellum, Cortex, Frontal Cortex BA9, Hippocampus, Hypothalamus, Nucleus accumbens basal ganglia, Putamen basal ganglia, and Substantia nigra) downloaded from https://www.gtexportal.org; Supplementary Note 1^29^). Of these, only SNPs overlapping open chromatin regions of brain-related samples (see previous paragraph) were included.

For 3D chromatin interaction mapping (Hi-C), we downloaded long-range chromatin interaction data from the adult cortex^24^ and human developing brain^30^ (Interactions to TSS for cortical plate and germinal zone, Tables S22 and S23 of Won *et al*.^30^). The enhancer region of these enhancer-promoter interactions was intersected with brain enhancers (see above) to only keep enhancer-promoter interactions overlapping known active brain enhancers. Then, the promoter region of these filtered enhancer-promoter interactions was mapped to a gene if it intersected with the region 250bp upstream and 500bp downstream of the corresponding gene transcription start site. SNPs were mapped to a gene if they overlapped the promoter of the filtered enhancer-promoter sites.

Finally, SNPs were positionally mapped to the nearest gene if the shortest distance to either transcription start site or end site was 60kb. This cutoff was selected because it maps the majority (90%) of SNPs to their nearest gene, following a distance distribution analysis.

The order of SNP-gene mapping was as follows: SNPs that mapped to a gene via brain eQTL or Hi-C interactions were prioritized and not also positionally mapped to a gene. A SNP was allowed to map to genes using both eQTL and Hi-C. SNPs without eQTL or Hi-C mappings were positionally mapped to a gene. Where a SNP positionally mapped to multiple genes, all associations were retained. These SNP-gene mappings were used for the pathway and gene set enrichment analysis described below, as well as to annotate SNPs from the GWAS analysis.

Using these criteria, 7.7% of SNPs mapped to genes using non-positional information (246,357 by eQTL and 16,923 by HiC, for a total of 263,280 SNPs); 2,917,948 SNPs mapped solely by positional information (89.2%). In total, SNPs mapped to 18,782 genes. 1,711,969 SNPs did not map to any genes (34.9%).

### Gene set enrichment analysis

For each of the nine CNB phenotypes, gene set enrichment analysis was performed using an implementation of GSEA for genetic variants^31,32^. GSEA was selected as it computes pathway enrichment scores using all available SNP information, which improves sensitivity, rather than using a hypergeometric model limited to SNPs passing a specific GWAS p-value cutoff. Moreover, pathway significance is ascertained using sample permutation, which corrects false-positives arising due to mapping of a few high-ranking SNPs to multiple nearby genes in the same pathway^33^. All SNPs were mapped to genes (as described in the “SNP-gene mapping for annotation and enrichment analyses” section above) and the gene score was defined as the best GWAS marginal p-value of all mapped SNPs for each gene. For each pathway, GSEA computes an enrichment score (ES) using the rank-sum of gene scores. The set of genes that appear in the ranked list before the rank-sum reaches its maximum deviation from zero, is called the “leading edge subset”, and is interpreted as the core set of genes responsible for the pathway’s enrichment signal. Following computation of the ES, we created a null distribution for each pathway by repeating genome-wide association tests with randomly label-permuted data and by computing ES from these permuted data; in this work, we use 100 permutations to reduce computational burden. As a test of sensitivity to this parameter, we increased this value to 1000 for the working memory phenotype (lnb_tp2). Finally, the ES on the original data is normalized to the score computed for the same gene set for label-permuted data (Z-score of real ES relative to mean of ES in label-permuted data), resulting in a Normalized Enrichment Score (NES) per pathway. The nominal p-value for the NES score is computed based on the null distribution and FDR correction is used to generate a q-value.

We used enrichment analysis to perform pathway analysis using pathway information compiled from HumanCyc^34^ (http://humancyc.org)^35^, NetPath (http://www.netpath.org)^36^., Reactome (http://www.reactome.org)., NCI Curated Pathways^37^, mSigDB^38^ (http://software.broadinstitute.org/gsea/msigdb/), and Panther^39^ (http://pantherdb.org/) and Gene Ontology^40^ (Human_GOBP_AllPathways_no_GO_iea_May_01_2018_symbol.gmt, downloaded from http://download.baderlab.org/EM_Genesets/May_01_2018/Human/symbol/Human_GOBP_AllPathways_no_GO_iea_May_01_2018_symbol.gmt); only pathways with 20-500 genes were used.

We also used enrichment analysis to perform a brain system and disease analysis using brain-related gene sets we compiled from various literature sources (see Supplementary Table 5 and Supplementary Note 5). Brain system gene sets included those identified through transcriptomic or proteomic assays in human brain tissue (i.e. direct measurement of expression), and genes associated with brain function by indirect inference (e.g. genetic association of nervous system disorders); both groups of gene sets were combined for this enrichment analysis. The transcriptomic/proteomic gene sets included: genes identified as markers for adult and fetal brain cell types using single-cell transcriptomic experiments^41-43^, genes enriched for brain-specific expression (Human Protein Atlas project (https://www.proteinatlas.org^44^); genes co-expressed with markers of various stages of human brain development (BrainSpan^45^); and genes encoding proteins altered in the schizophrenia synaptosomal proteome^46^. Brain disease gene sets included: genes associated with schizophrenia, bipolar disorder, autism spectrum disorder and major depressive disorder through large-scale genetic association studies by the Psychiatric Genomics Consortium (Supplementary Note 5); genes associated with nervous system disorders by the Human Phenotype Ontology^47^. Genes in the second group were filtered to only include genes with detectable expression in the fetal^48^ or adult human brain^44^. A total of 1,321 gene sets were collected across both system and disease categories (Table S14). Only gene sets with 20-500 genes were included in the analysis; 421 gene sets met these criteria and were included in the enrichment analysis.

### Leading edge gene interaction network

Genes contributing to pathway enrichment results (leading edge genes) were obtained in our GSEA analysis for genetic variants^31^. A gene-gene interaction network was constructed from leading edge genes of pathways with q < 0.05 using the online GeneMANIA service (v 3.6.0; https://genemania.org^49^.) (human database, default settings); the resulting network and edge attributes were downloaded. This network was imported into Cytoscape v3.7.1 for visualization. Known drug associations were obtained from DGIdb^50^ and GWAS associations with nervous system disorders were obtained from the NHGRI-EBI GWAS catalogue, via programmatic search using the TargetValidation.org API^51,52^. Cell type marker information was compiled from single cell RNA-seq datasets, including those for adult and fetal human brain^41-43^.

## Results

We developed a systems-genomics analysis workflow to identify genetic variants associated with normal cognitive phenotypes (Figure 1). Briefly, genotypes were imputed using a reference panel from the 1,000 Genomes Project^53^, and samples were limited to those of European genetic ancestry (Supplementary Figure 1-3, Supplementary Table 1). 3,394 individuals and ∼4.9M SNPs passed the quality control and imputation process. Following quality control of phenotype data, 3,116 European samples passed both genotype and phenotype filters and were included in downstream analyses. We selected nine phenotypes from the Penn Computerized Neurocognitive Test Battery (CNB) representing overall accuracy in four cognitive domains: complex cognition, executive function, declarative memory, and social processing (Supplementary Table 3). Measures included performance for verbal reasoning, nonverbal reasoning, spatial reasoning, attention allocation, working memory, recall tests for faces, words and objects, and emotion identification^11^. In all instances, age and sex was regressed out of the phenotype (Supplementary Table 4) and samples were thereafter binarized into poor and good performers (bottom and top 33% percentile, respectively) resulting in ∼1,000 samples per group for each phenotype (Supplementary Figure 4,5, Supplementary Table 3).

**Figure 1.**
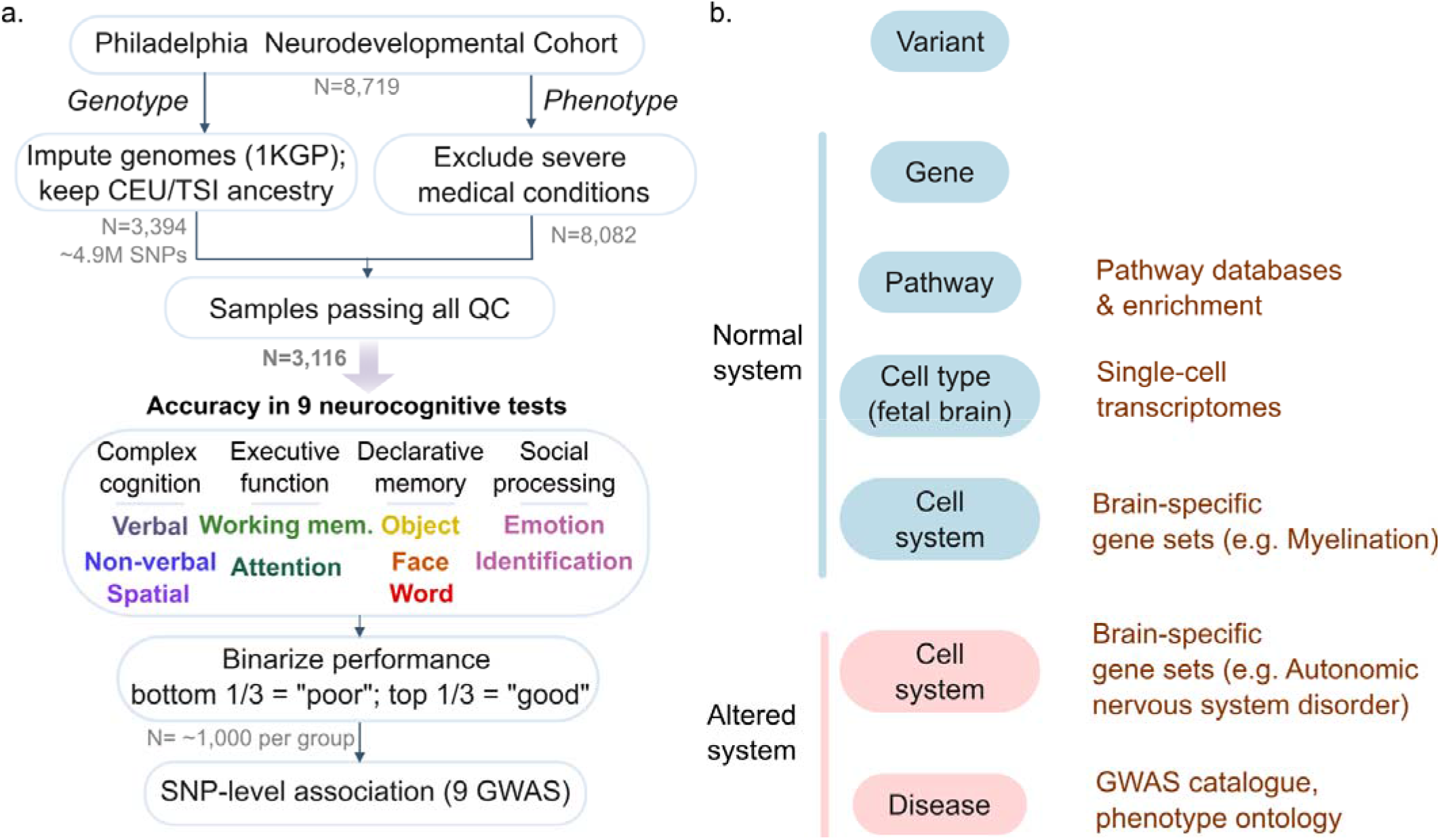
Framework for multi-scale systems-genomics analysis for neurocognitive phenotypes from the Philadelphia Neurodevelopmental Cohort. a. Workflow for genome-wide association analysis (GWAS). Genotypes were imputed (1KGP reference), and limited to European samples. Samples with severe medical conditions were removed and invalid test scores excluded. Nine neurocognitive test scores were binarized after regressing out age and sex. GWAS was performed using the accuracy measure as a phenotype for each of these nine phenotypes. b. Framework to organize variant-level associations into a multi-scale systems view in health (blue) and disease (red). Existing functional genomic resources used for annotation shown in brown.

For each of the nine phenotypes, we first performed SNP-level genome-wide association analysis, as a comparative baseline following traditional methods. We used a mixed-effects linear model that included genome-wide genetic ancestry as a covariate (GCTA^22^). Among the nine phenotypes, 661 SNPs had suggestive levels of significance at the genome-wide level (p < 10^−5^; Figure 1b,c, Supplementary Figure 6,7,8, Supplementary Table 6). Over half of these SNPs are associated with tasks related to complex cognition, i.e. verbal reasoning, non-verbal reasoning and spatial reasoning (377 SNPs or 57%). 27% were associated with executive function (177 SNPs), which included attention allocation and working memory. 13% SNPs were associated with declarative memory tasks (83 SNPs), which included face recall, word recall and object recall. 4% of SNPs were associated with emotion identification (24 SNPs), a measure of social processing. More generally, SNPs associated with PNC cognitive phenotypes at suggestive significance levels (p<10^−5^) map to genes previously associated with diseases of the nervous system and/or mark cell-types in the fetal and newborn brain^41,43^ (Supplementary Figure 8, Supplementary Table 7). We predict that one-sixth of suggestive peaks (112 SNPs or 17%) are linked to a functional consequence in brain tissue, including non-synonymous changes to protein sequence (Supplementary Fig. 8), presence in brain-specific promoters and enhancers, or association with changes in gene expression (Supplementary Table 6).

Nonverbal reasoning was the only phenotype with SNPs passing the cutoff for genome-wide significance (rs77601382 and rs5765534, p = 4.6×10^−8^) (Figure 2). The peak is located in a ∼33kb region (chr22:45,977,415-46,008,175) overlapping the 3’ end of the Fibulin-1 (*FBLN1*) gene, including the last intron and exon (Figure 2b). To better understand the significance of this gene in brain function, we examined *FBLN1* expression in published fetal and adult transcriptomes, and single-cell data^29,43,45^. *FBLN1* transcription in the human brain is highest in the early stages of fetal brain development, with little to no expression in the adult (Figure 2c, Supplementary Figure 8); this is consistent with single-cell assays showing *FBLN1* to be a marker for dividing progenitor cells in the fetal brain^43^. *FBLN1* encodes a glycoprotein present in the extracellular matrix; this protein is a direct interactor of proteins involved in neuronal diseases, such as Amyloid Precursor Protein-1^54^ (Supplementary Figure 9 ^55^). *FBLN1* expression is upregulated in the brain in schizophrenia and has been previously associated with genetic risk for bipolar disorder (Figure 1d, ^56,57^). Therefore, we conclude that *FBLN1*, associated with nonverbal reasoning test performance, shows characteristics of a gene involved in neurodevelopment and the dysregulation of which could increase risk for psychotic disorders of neurodevelopmental origin.

**Figure 2.**
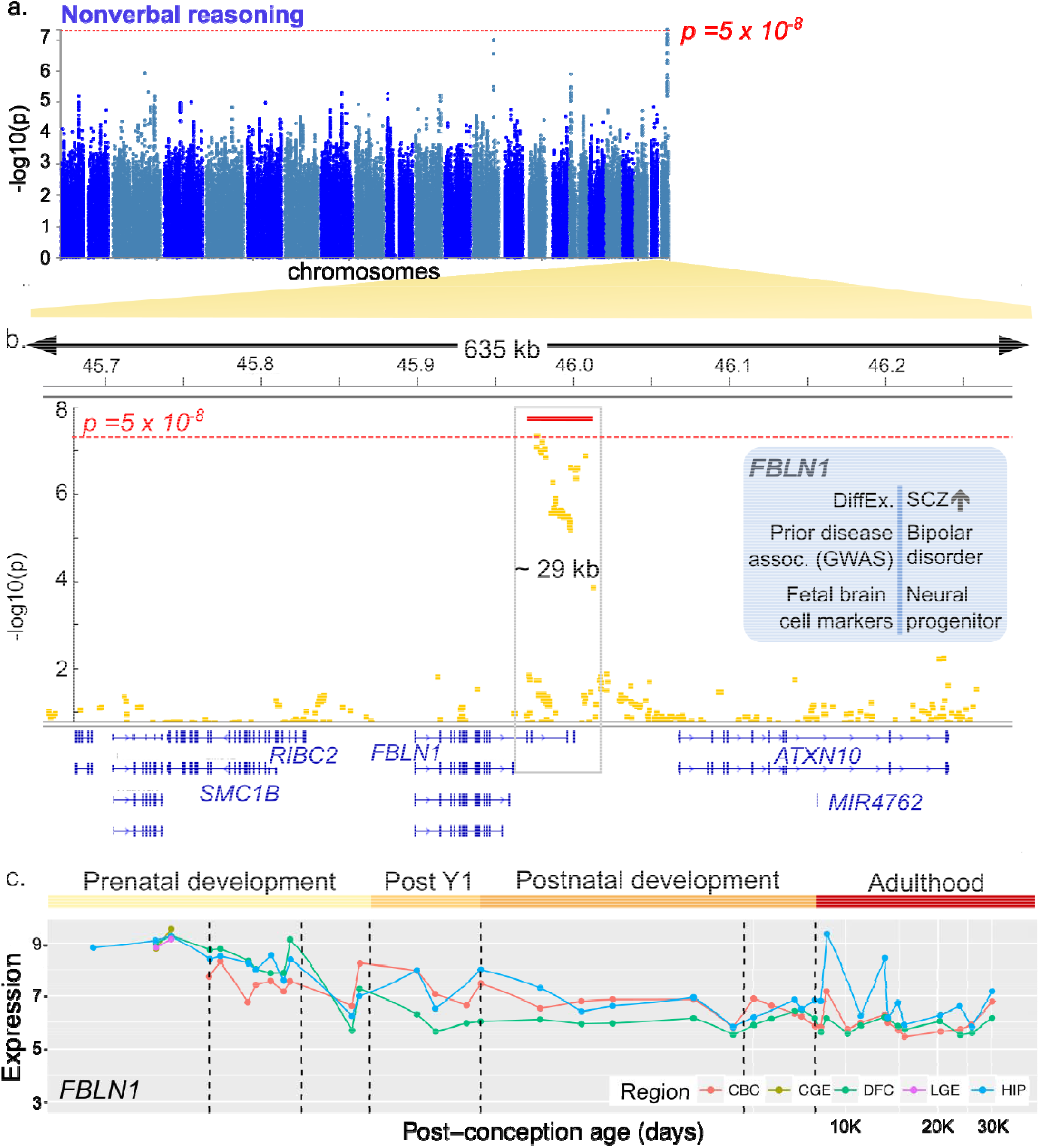
Genome-wide significance of *FBLN1* region for binarized performance in nonverbal reasoning a. Manhattan plot of univariate SNP association with binarized performance in nonverbal reasoning (N=1,024 poor vs. 1,023 good performers; 4,893,197 SNPs). Plot generated using FUMA^70^. b. Detailed view of hit region at chr22q13. Two SNPs pass genome-wide significance threshold, rs77601382 and rs74825248 (p=4.64e-8). View using Integrated Genome Viewer (v2.3.93^71,72^). The red bar indicates the region with increased SNP-level association. c. *FBLN1* transcription in the human brain through the lifespan. Data from BrainSpan^45^. Log-transformed normalized expression is shown for cerebellar cortex (CBC), central ganglionic eminence (CGE) and lateral ganglionic eminence (LGE), dorsal frontal cortex (DFC), and hippocampus (HIP).

Pathway analysis is an established systems-genomics technique used to improve the statistical power of subthreshold univariate signal by aggregation of signal and reduction of multiple hypothesis testing burden, as well as to provide mechanistic insight into cellular processes that affect phenotypic outcome. Pathway analysis has been successfully used to link genetic disease risk to cellular processes for diseases in various domains, including schizophrenia^58^, breast cancer^59^ and type 2 diabetes^60^. We performed pathway analysis for the nine phenotypes using a rank-based pathway analysis strategy (GSEA^31,38^, 500 permutations; 4,102 pathways tested). SNPs were mapped to genes using brain-specific eQTL, chromatin interaction and positional information, using the same method as described above. The working memory phenotype demonstrated significant enrichment of top-ranking genetic variants in a developmental pathway (q < 0.05; Supplementary Tables 8-10), showing biologically relevant signal where our univariate SNP-based baseline analysis did not. An advantage of the rank-based pathway analysis over those based on hypergeometric or binomial tests, is that the former provides a list of “leading-edge” genes driving the pathway-level enrichment signal, which can be further interpreted. We annotated leading-edge genes with prior knowledge about genetic associations with nervous system disorders, transcription in brain cell types^41-43,51^ and known drug interactions^50^. Out of 53 leading edge genes of this gene set, roughly one-half are known brain cell markers (25 genes or 47%), roughly one-third have known drug interactions (17 genes or 36%), and ∼11% are associated with nervous system disease (6 genes) (pathway q < 0.05, Figure 3a, Supplementary Table 10, Supplementary Figure 11). Among disease-associated genes were those associated with autism (*CSDE1*) and Parkinson’s disease (*LHFPL2*).

**Figure 3.**
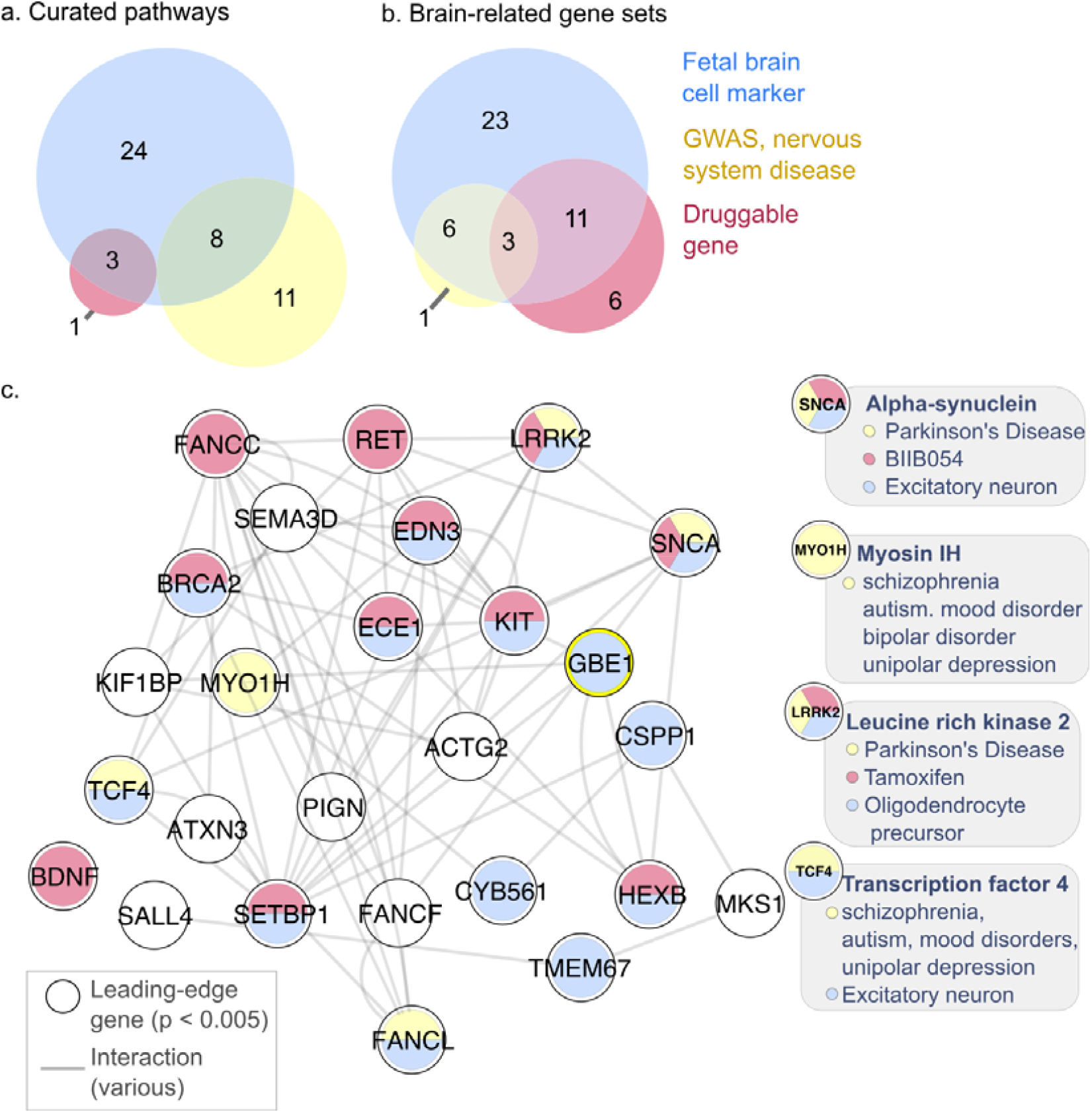
Pathway and brain system and disease analysis for the working memory performance phenotype a. Attributes of leading edge genes in pathway gene sets associated (q < 0.05) with working memory. Colours indicate transcription in brain cell types (blue), genetic associations with nervous system disorders (yellow), or those with known drug targets (pink) (N=53 genes in total; 47 with annotations). b. Leading edge genes in brain-related gene sets associated with disease, drugs or brain cell types (N=71 genes total; 50 with annotations). Details in Supplementary Table. Legend as in a. c. Gene-gene interaction network for working memory leading edge genes from enriched (q < 0.05) brain-related gene sets. Only genes with top SNP p < 5×10^−3^ are shown (26 genes). Nodes show genes and fill colour indicates genes associated with brain cell types, drugs or genetic associations with nervous system disorders (colours as in panel a, white indicates absence of association). Edges indicate known interactions (from GeneMANIA^49^). Genes from the network with disease associations are highlighted with grey description bubbles.

To perform a brain system and disease analysis, we performed a second enrichment analysis using gene sets curated from the literature, including transcriptomic and proteomic profiles of the developing and adult healthy brain and brains affected by mental illness, brain-related genome-wide association studies, and terms from a phenotype ontology (421 gene sets tested, Supplementary Note 5, Supplementary Table 5, Supplementary Data 1). Two gene sets pertaining to general nervous system dysfunction were significantly enriched (q<0.05; GSEA, 500 permutations), again related to working memory (Figure 3c, Supplementary Table 11). Roughly 17% of the 71 leading edge genes of these gene sets are associated with nervous system disorders (12 genes), roughly one-third have predicted drug targets (22 genes, 31%), and over half (43 genes or 61%) are markers of brain cell-types (Figure 3b,c; Supplementary Table 12, 13). Two genes have all three attributes: *SNCA* and *LRRK2* (Figure 3c, Supplementary Table 13). Leading edge genes have genetic associations, including those with schizophrenia, autism spectrum disorder, Parkinson’s disease, Alzheimer’s disease, depression and mood disorders (Figure 3c, Supplementary Table 13). In summary, we identified many genetic variants associated with normative variation in a range of neurocognitive phenotypes enriched in pathways and gene sets related to development, nervous system dysfunction and mental disorders.

## Discussion

To our knowledge, this is the first study to identify genetic variants that may contribute to normal human variation in multiple, diverse cognitive domains, and to link these to various levels of brain system organization, including genes, pathways, cell types, brain regions, diseases and known drug targets. These associations, particularly potential drug targets, represent hypotheses to be experimentally validated in model systems to improve the mechanistic understanding of the molecular substrates of the respective phenotypes.

We found an enrichment of genetic variants associated with complex cognitive phenotypes (75-219 suggestive peaks in a Manhattan plot), consistent with heritability estimates of up to 0.30-0.41 for these phenotypes^12^. We also found that many variants, genes and pathways associated with normal variation in neurocognitive phenotypes have known roles in neurodevelopment, modulating gene expression in the fetal and adult brain and increasing risk for psychiatric diseases of neurodevelopmental origin (Figure 1, Supplementary Table 6, 7, 10, 13). Multiple lines of evidence suggest that *FBLN1*, the gene we find associated with genome-wide significant SNPs for nonverbal reasoning, is dysregulated in brain-related disease. In addition to the evidence provided in our results (Figure 2c, Supplementary Figure 8,9), the *FBLN1* gene has been associated with a rare genetic syndrome that includes various cognitive impairments, and protein levels of *FBLN1* have been associated with altered risk for ischaemic stroke^61,62^. However, the mechanism by which *FBLN1* contributes to normal brain function is not known. We also do not exclude the possibility that suggestive peaks we identified within *FBLN1* may affect the function of neighbouring or otherwise linked genes, which may instead or in combination affect the phenotype. One such gene is Ataxin-10 (*ATXN10*), which is the next neighboring downstream gene to *FBLN1*, in which a pentanucleotide repeat expansion causes spinocerebellar atrophy and ataxia^63^. The *FBLN1* locus was not significantly enriched in a large GWAS study of general cognitive ability ^64^, suggesting that this locus may be influencing a specialized trait.

A limitation of the current study is the relatively small size of the patient cohort – roughly 1,000 cases and controls each per phenotype – compared to contemporary GWAS studies which may include over 100,000 individuals. The reduced sample size is partly because we chose to limit the analysis to individuals with European genetic ancestry, to maintain the largest number of samples while avoiding the confound with genetic ancestry. Furthermore, we dichotomized the phenotype into bottom and top performers, ignoring samples in the middle, as our goal was to work with a subset enriched for extremes within typical phenotypic variation, to strengthen signal. For all phenotypes tested in this work, we also performed genome-wide association tests using continuously-valued measures, instead of binarized phenotypes; none of the associations resulted in significant results (data not shown). This lack of association is consistent with the strategy to binarize outcomes for improved contrast; binarization includes only the top and bottom thirds of performance measures, and ignores the measures in between.

This work contributes towards an understanding of the molecular and systems-level underpinnings of individual cognitive tasks. These associations will need to be validated in better-powered datasets, possibly using newer neurobehavioural measurement standards in the field^65^ but can currently be used as hypotheses to plan biological experiments, or as support for orthogonal methods studying the relevance of genes and pathways we identify for brain biology. Studying the overlap in genetic architecture between these phenotypes, similar to cross-disorder genetic studies^66^, may also inform disease classification^67,68^. Our analysis is limited to univariate genetic effects, and future work should explore the contribution of interactions between individual SNPs^69^, though this will require many more samples per phenotype. We propose that research frameworks for linking genotype to phenotype for brain-related traits include systems genomics analysis, considering pathways, cells, anatomical structures, and physiological processes as organizational layers to improve the amount of genetic signal that can be extracted from available genetic data, which otherwise would be missed if just considering SNPs and genes. For example, the working memory phenotype had no significant SNPs that met the genome-wide significance cutoff. However, gene sets related to development and autonomic nervous system dysfunction demonstrated significant clustering of high-ranking variants, including those in *SLIT3* (rs62376937) and *ROBO2* (rs12497629), which mediate axon guidance in the developing nervous system. The conceptual strategy we outline in this work, of organizing variant-related annotation into a systems-level view is generalizable across biomedical domains and to human disease (Figure 1,4). Integration of such evidence across studies can identify common themes or discrepancies to encourage thinking of a systems-level view of genotype-phenotype association for disease.

**Figure 4.**
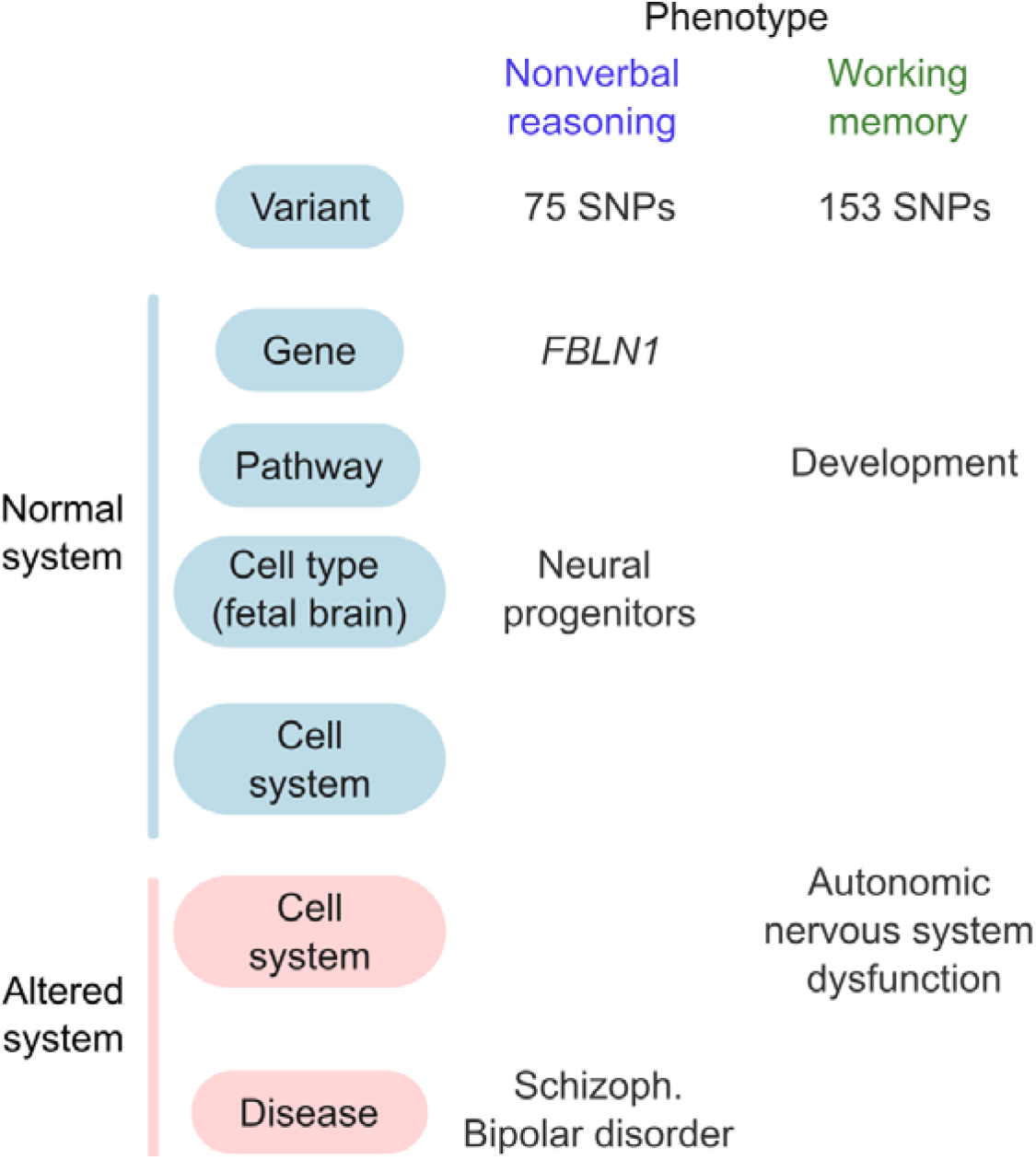
Summary of evidence linking genetic variants associated with cognitive test performance to multiple levels of brain organization. Each column shows data for an individual phenotype, grou ed by phenotype domain; rows show associations at increasingly more general scales (from top to bottom); evidence linking variants to healthy system and disease system shown in blue and red, respectively. Circles indicate relative number of suggestive variant peaks (p < 10^−5^) from GWAS). Pathways and cell systems are those identified by gene set enrichment analyses (q<0.05). Cell types are those for which *FBLN1* is found to be a marker from single-cell transcriptome data^43^. Gene-disease associations are identified for significant SNPs, using gene-disease mappings from the NHGRI-EBI catalogue^51^.

## References

1 Lett, T. A., Voineskos, A. N., Kennedy, J. L., Levine, B. & Daskalakis, Z. J. Treating working memory deficits in schizophrenia: a review of the neurobiology. Biological psychiatry 75, 361–370, doi:10.1016/j.biopsych.2013.07.026 (2014).

2 Insel, T. et al. Research domain criteria (RDoC): toward a new classification framework for research on mental disorders. (2010).

3 Hill, W. D. et al. A combined analysis of genetically correlated traits identifies 187 loci and a role for neurogenesis and myelination in intelligence. Molecular psychiatry 24, 169–181, doi:10.1038/s41380-017-0001-5 (2019).

4 Okbay, A. et al. Genome-wide association study identifies 74 loci associated with educational attainment. Nature 533, 539–542, doi:10.1038/nature17671 (2016).

5 Sniekers, S. et al. Genome-wide association meta-analysis of 78,308 individuals identifies new loci and genes influencing human intelligence. Nature genetics 49, 1107-1112, doi:10.1038/ng.3869 (2017).

6 Blokland, G. A. et al. Quantifying the heritability of task-related brain activation and performance during the N-back working memory task: a twin fMRI study. Biol Psychol 79, 70–79, doi:10.1016/j.biopsycho.2008.03.006 (2008).

7 Braver, T. S. et al. A parametric study of prefrontal cortex involvement in human working memory. NeuroImage 5, 49–62, doi:10.1006/nimg.1996.0247 (1997).

8 Minzenberg, M. J., Laird, A. R., Thelen, S., Carter, C. S. & Glahn, D. C. Meta-analysis of 41 functional neuroimaging studies of executive function in schizophrenia. Arch Gen Psychiatry 66, 811–822, doi:10.1001/archgenpsychiatry.2009.91 (2009).

9 Calkins, M. E. et al. The Philadelphia Neurodevelopmental Cohort: constructing a deep phenotyping collaborative. Journal of child psychology and psychiatry, and allied disciplines 56, 1356–1369, doi:10.1111/jcpp.12416 (2015).

10 Satterthwaite, T. D. et al. Neuroimaging of the Philadelphia neurodevelopmental cohort. NeuroImage 86, 544–553, doi:10.1016/j.neuroimage.2013.07.064 (2014).

11 Gur, R. C. et al. A cognitive neuroscience-based computerized battery for efficient measurement of individual differences: standardization and initial construct validation. Journal of neuroscience methods 187, 254–262, doi:10.1016/j.jneumeth.2009.11.017 (2010).

12 Robinson, E. B. et al. The genetic architecture of pediatric cognitive abilities in the Philadelphia Neurodevelopmental Cohort. Molecular psychiatry 20, 454–458, doi:10.1038/mp.2014.65 (2015).

13 Germine, L. et al. Association between polygenic risk for schizophrenia, neurocognition and social cognition across development. Translational psychiatry 6, e924, doi:10.1038/tp.2016.147 (2016).

14 Luna, B., Garver, K. E., Urban, T. A., Lazar, N. A. & Sweeney, J. A. Maturation of cognitive processes from late childhood to adulthood. Child development 75, 1357–1372, doi:10.1111/j.1467-8624.2004.00745.x (2004).

15 Simmonds, D. J., Hallquist, M. N. & Luna, B. Protracted development of executive and mnemonic brain systems underlying working memory in adolescence: A longitudinal fMRI study. NeuroImage 157, 695–704, doi:10.1016/j.neuroimage.2017.01.016 (2017).

16 Glessner, J. T. et al. Strong synaptic transmission impact by copy number variations in schizophrenia. Proceedings of the National Academy of Sciences of the United States of America 107, 10584–10589, doi:10.1073/pnas.1000274107 (2010).

17 Verma, S. S. et al. Imputation and quality control steps for combining multiple genome-wide datasets. Frontiers in Genetics 5, 370 (2014).

18 Hinrichs, A. S. et al. The UCSC Genome Browser Database: update 2006. Nucleic acids research 34, D590–598, doi:10.1093/nar/gkj144 (2006).

19 Delaneau, O., Zagury, J. F. & Marchini, J. Improved whole-chromosome phasing for disease and population genetic studies. Nature methods 10, 5–6, doi:10.1038/nmeth.2307 (2013).

20 Howie, B. N., Donnelly, P. & Marchini, J. A flexible and accurate genotype imputation method for the next generation of genome-wide association studies. PLoS genetics 5, e1000529, doi:10.1371/journal.pgen.1000529 (2009).

21 Anderson, C. A. et al. Data quality control in genetic case-control association studies. Nature protocols 5, 1564–1573, doi:10.1038/nprot.2010.116 (2010).

22 Yang, J., Lee, S. H., Goddard, M. E. & Visscher, P. M. GCTA: a tool for genome-wide complex trait analysis. American journal of human genetics 88, 76–82, doi:10.1016/j.ajhg.2010.11.011 (2011).

23 Schmidt, A. et al. Acute effects of heroin on negative emotional processing: relation of amygdala activity and stress-related responses. Biological psychiatry 76, 289–296, doi:10.1016/j.biopsych.2013.10.019 (2014).

24 Schmitt, A. D. et al. A Compendium of Chromatin Contact Maps Reveals Spatially Active Regions in the Human Genome. Cell reports 17, 2042–2059, doi:10.1016/j.celrep.2016.10.061 (2016).

25 Krueger F. Trim Galore!, <http://www.bioinformatics.babraham.ac.uk/projects/trim_galore/> (

26 Wingett, S. et al. HiCUP: pipeline for mapping and processing Hi-C data. F1000Research 4, 1310, doi:10.12688/f1000research.7334.1 (2015).

27 Mifsud, B. et al. GOTHiC, a probabilistic model to resolve complex biases and to identify real interactions in Hi-C data. PloS one 12, e0174744, doi:10.1371/journal.pone.0174744 (2017).

28 Kundaje, A. et al. Integrative analysis of 111 reference human epigenomes. Nature 518, 317–330, doi:10.1038/nature14248 (2015).

29 Battle, A., Brown, C. D., Engelhardt, B. E. & Montgomery, S. B. Genetic effects on gene expression across human tissues. Nature 550, 204–213, doi:10.1038/nature24277 (2017).

30 Won, H. et al. Chromosome conformation elucidates regulatory relationships in developing human brain. Nature 538, 523–527, doi:10.1038/nature19847 (2016).

31 Wang, K., Li, M. & Bucan, M. Pathway-based approaches for analysis of genomewide association studies. American journal of human genetics 81, 1278–1283, doi:10.1086/522374 (2007).

32 Wang, K., Li, M. & Hakonarson, H. Analysing biological pathways in genome-wide association studies. Nature reviews. Genetics 11, 843–854, doi:10.1038/nrg2884 (2010).

33 Mooney, M. A., Nigg, J. T., McWeeney, S. K. & Wilmot, B. Functional and genomic context in pathway analysis of GWAS data. Trends Genet 30, 390–400, doi:10.1016/j.tig.2014.07.004 (2014).

34 Romero, P. et al. Computational prediction of human metabolic pathways from the complete human genome. Genome biology 6, R2, doi:10.1186/gb-2004-6-1-r2 (2005).

35 Kandasamy, K. et al. NetPath: a public resource of curated signal transduction pathways. Genome biology 11, R3, doi:10.1186/gb-2010-11-1-r3 (2010).

36 Fabregat, A. et al. The Reactome pathway Knowledgebase. Nuc Acids Res 44, D481–487, doi:10.1093/nar/gkv1351 (2016).

37 Schaefer, C. F. et al. PID: the Pathway Interaction Database. Nucleic acids research 37, D674–679, doi:10.1093/nar/gkn653 (2009).

38 Subramanian, A. et al. Gene set enrichment analysis: a knowledge-based approach for interpreting genome-wide expression profiles. Proceedings of the National Academy of Sciences of the United States of America 102, 15545–15550, doi:10.1073/pnas.0506580102 (2005).

39 Mi, H. et al. The PANTHER database of protein families, subfamilies, functions and pathways. Nucleic acids research 33, D284–288, doi:10.1093/nar/gki078 (2005).

40 Consortium, T. G. O. The Gene Ontology Resource: 20 years and still GOing strong. Nucleic acids research 47, D330–d338, doi:10.1093/nar/gky1055 (2019).

41 Darmanis, S. et al. A survey of human brain transcriptome diversity at the single cell level. Proceedings of the National Academy of Sciences of the United States of America 112, 7285–7290, doi:10.1073/pnas.1507125112 (2015).

42 Lake, B. B. et al. Neuronal subtypes and diversity revealed by single-nucleus RNA sequencing of the human brain. Science (New York, N.Y.) 352, 1586–1590, doi:10.1126/science.aaf1204 (2016).

43 Nowakowski, T. J. et al. Spatiotemporal gene expression trajectories reveal developmental hierarchies of the human cortex. Science (New York, N.Y.) 358, 1318–1323, doi:10.1126/science.aap8809 (2017).

44 Yu, N. Y. et al. Complementing tissue characterization by integrating transcriptome profiling from the Human Protein Atlas and from the FANTOM5 consortium. Nucleic acids research 43, 6787–6798, doi:10.1093/nar/gkv608 (2015).

45 Kang, H. J. et al. Spatio-temporal transcriptome of the human brain. Nature 478, 483–489, doi:10.1038/nature10523 (2011).

46 Velasquez, E. et al. Synaptosomal Proteome of the Orbitofrontal Cortex from Schizophrenia Patients Using Quantitative Label-Free and iTRAQ-Based Shotgun Proteomics. Journal of proteome research 16, 4481–4494, doi:10.1021/acs.jproteome.7b00422 (2017).

47 Kohler, S. et al. Expansion of the Human Phenotype Ontology (HPO) knowledge base and resources. Nucleic acids research 47, D1018–d1027, doi:10.1093/nar/gky1105 (2019).

48 Zhong, S. et al. A single-cell RNA-seq survey of the developmental landscape of the human prefrontal cortex. Nature 555, 524–528, doi:10.1038/nature25980 (2018).

49 Franz, M. et al. GeneMANIA update 2018. Nucleic acids research 46, W60–w64, doi:10.1093/nar/gky311 (2018).

50 Cotto, K. C. et al. DGIdb 3.0: a redesign and expansion of the drug-gene interaction database. Nucleic acids research 46, D1068–d1073, doi:10.1093/nar/gkx1143 (2018).

51 Buniello, A. et al. The NHGRI-EBI GWAS Catalog of published genome-wide association studies, targeted arrays and summary statistics 2019. Nucleic acids research 47, D1005–d1012, doi:10.1093/nar/gky1120 (2019).

52 Carvalho-Silva, D. et al. Open Targets Platform: new developments and updates two years on. Nucleic acids research 47, D1056–d1065, doi:10.1093/nar/gky1133 (2019).

53 1000 Genomes Project Consortium et al. A global reference for human genetic variation. Nature 526, 68–74, doi:10.1038/nature15393 (2015).

54 Ohsawa, I., Takamura, C. & Kohsaka, S. Fibulin-1 binds the amino-terminal head of beta-amyloid precursor protein and modulates its physiological function. J Neurochem 76, 1411–1420, doi:10.1046/j.1471-4159.2001.00144.x (2001).

55 Stark, C. et al. BioGRID: a general repository for interaction datasets. Nucleic acids research 34, D535–539, doi:10.1093/nar/gkj109 (2006).

56 Wang, D. et al. Comprehensive functional genomic resource and integrative model for the human brain. Science (New York, N.Y.) 362, doi:10.1126/science.aat8464 (2018).

57 Greenwood, T. A., Akiskal, H. S., Akiskal, K. K. & Kelsoe, J. R. Genome-wide association study of temperament in bipolar disorder reveals significant associations with three novel Loci. Biological psychiatry 72, 303–310, doi:10.1016/j.biopsych.2012.01.018 (2012).

58 Biological insights from 108 schizophrenia-associated genetic loci. Nature 511, 421–427, doi:10.1038/nature13595 (2014).

59 Michailidou, K. et al. Association analysis identifies 65 new breast cancer risk loci. Nature 551, 92–94, doi:10.1038/nature24284 (2017).

60 Xue, A. et al. Genome-wide association analyses identify 143 risk variants and putative regulatory mechanisms for type 2 diabetes. Nature communications 9, 2941, doi:10.1038/s41467-018-04951-w (2018).

61 Palumbo, P. et al. Clinical and molecular characterization of an emerging chromosome 22q13.31 microdeletion syndrome. American journal of medical genetics. Part A 176, 391–398, doi:10.1002/ajmg.a.38559 (2018).

62 Vadgama, N., Lamont, D., Hardy, J., Nasir, J. & Lovering, R. C. Distinct proteomic profiles in monozygotic twins discordant for ischaemic stroke. Molecular and cellular biochemistry, doi:10.1007/s11010-019-03501-2 (2019).

63 Matsuura, T. et al. Large expansion of the ATTCT pentanucleotide repeat in spinocerebellar ataxia type 10. Nature genetics 26, 191–194, doi:10.1038/79911 (2000).

64 Savage, J. E. et al. Genome-wide association meta-analysis in 269,867 individuals identifies new genetic and functional links to intelligence. Nature genetics 50, 912–919, doi:10.1038/s41588-018-0152-6 (2018).

65 Weintraub, S. et al. Cognition assessment using the NIH Toolbox. Neurology 80, S54–64, doi:10.1212/WNL.0b013e3182872ded (2013).

66 Lee, S. H. et al. Genetic relationship between five psychiatric disorders estimated from genome-wide SNPs. Nature genetics 45, 984–994, doi:10.1038/ng.2711 (2013).

67 Clementz, B. A. et al. Identification of Distinct Psychosis Biotypes Using Brain-Based Biomarkers. The American journal of psychiatry 173, 373–384, doi:10.1176/appi.ajp.2015.14091200 (2016).

68 Jeste, S. S. & Geschwind, D. H. Disentangling the heterogeneity of autism spectrum disorder through genetic findings. Nature reviews. Neurology 10, 74–81, doi:10.1038/nrneurol.2013.278 (2014).

69 Wang, W. et al. Pathway-based discovery of genetic interactions in breast cancer. PLoS genetics 13, e1006973, doi:10.1371/journal.pgen.1006973 (2017).

70 Watanabe, K., Taskesen, E., van Bochoven, A. & Posthuma, D. Functional mapping and annotation of genetic associations with FUMA. Nature communications 8, 1826, doi:10.1038/s41467-017-01261-5 (2017).

71 Robinson, J. T. et al. Integrative genomics viewer. Nature biotechnology 29, 24–26, doi:10.1038/nbt.1754 (2011).

72 Thorvaldsdottir, H., Robinson, J. T. & Mesirov, J. P. Integrative Genomics Viewer (IGV): high-performance genomics data visualization and exploration. Briefings in bioinformatics 14, 178–192, doi:10.1093/bib/bbs017 (2013).

