## Supplementary Notes and Figures for "Multi-scale systems genomics analysis predicts pathways, cell types and drug targets involved in normative human cognition variation"

Genetics and Pathway Analysis of Normative Cognitive Variation in the Philadelphia Neurodevelopmental Cohort

### Supplementary Information

#### Supplementary Note 1: Variant annotation sources

| **Annotation type** | **Web source (if applicable)** | **Ref** |
| --- | --- | --- |
| Brain enhancers and promoters | <https://egg2.wustl.edu/roadmap/data/byFileType/chromhmmSegmentations/ChmmModels/core_K27ac/jointModel/final/all.mnemonics.bedFiles.tgz> from  <https://egg2.wustl.edu/roadmap/web_portal/chr_state_learning.html>  downloaded on 3 Jan 2019  Enhancer states = { 7_Enh, 6_EnhG} ; Promoter states = { TssA 2_TssAFlnk }; from E067 (angular gyrus), E071 (hippocampus middle), E072 (inferior temporal lobe), E073 (dorsolateral prefrontal cortex). | ^1^ |
| Brain eQTLs | GTEx: <https://storage.googleapis.com/gtex_analysis_v7/single_tissue_eqtl_data/GTEx_Analysis_v7_eQTL.tar.gz> from <https://www.gtexportal.org/home/datasets>  downloaded on 13 Nov 2017  Gene-variant pairs were obtained from the "signif_gene_variant_pairs" files. | ^2,3^ |
| Fetal brain eQTL | Supplementary Table 3 of main publication | ^4^ |
| Brain fQTL and multiQTL | <http://resource.psychencode.org/Datasets/Derived/QTLs/DER-11_hg19_fQTL.significant.txt> and <http://resource.psychencode.org/Datasets/Derived/QTLs/DER-12_hg19_multiQTL.list.txt>  From <http://resource.psychencode.org/#Integrative>  downloaded on 27 March 2019 | ^5^ |
| SNP functional consequence | Biomart: <https://www.ensembl.org/biomart/martview/ea36e1db44508c060447c5076ff923e0>  Database: Ensembl Variation 96  Dataset: Human Short Variants (SNPs and indels excluding flagged variants) (GRCh37.p13); Filter: By variant name; downloaded on 5 April 2019 | ^6^ |
| GWAS associations from literature | Systematic gene searches at TargetValidation.org, with filter for ontology code EFO_0000618 ("nervous system disorders") | ^7,8^ |

#### Supplementary Note 2: Gene annotation sources

| **Annotation type** | **Source** | **Ref** |
| --- | --- | --- |
| Differential expression in psychiatric disease | <https://www.synapse.org/#!Synapse:syn5607652>  downloaded from Synapse CommonMind Portal <https://www.synapse.org/#!Synapse:syn2759792> | ^3^ |
| GWAS associations from literature | Systematic gene searches at TargetValidation.org, with filter for ontology code EFO_0000618 ("nervous system disorders") | ^7,8^ |
| Fetal brain scRNA | From Supplementary Table 5 | ^9^ |
| Adult brain cell types | <http://resource.psychencode.org/Datasets/Derived/SC_Decomp/DER-19_Single_cell_markergenes_TPM.xlsx> | ^10,11^ |
| Gene-drug interactions | http://www.dgidb.org/data/interactions.tsv | ^12^ |
| Expression during human brain lifespan | Supplementary Table 12 | ^13^ |

#### Supplementary Note 3: SNP annotation method

Details of annotation sources are in Supplementary Note 1. SNPs were annotated by matching dbSNP identifiers or location. SNPs overlapping coordinates for brain enhancers or promoters were mapped to those categories. For the following categories, SNPs were matched by rsID: Brain fQTLs and multiQTL from PsychEncode^5^; fetal brain eQTLs^4^; SNP functional consequence from Biomart^6^; and GWAS associations^7^. GWAS associations from literature were filtered to include studies with sample size > 1000 and genome wide significance threshold of 5x10^-8^. Brain eQTLs from GTEx were matched using the "variant_id" column providing genomic location in the format "chr_position_allele1_allele2" while ignoring further suffixes.

#### Supplementary Note 4: Gene annotation method

Details of annotation sources are listed in Supplementary Note 2. GWAS-gene annotations were downloaded from TargetValidation.org^8^ using the Open Targets Platform REST API. Only associations for nervous system disorders were downloaded. Only those associations in GWAS studies with sample sizes greater than 1,000 individuals and with nominal p-value less than 5x10^-5^ were considered; genes were mapped to associated diseases even if the associated variant was different than that found in this study. This was done because the goal for this analysis was to identify genetic associations at the gene-level, rather than variant-level, with the disease. For all other annotation sources, genes were matched by name.

#### Supplementary Note 5: Compilation of brain-related gene sets

Brain-related datasets were broadly of two categories: those directly measuring gene activity in the brain, including transcriptomic and proteomics studies, and those inferred from analyses such as GWAS.

##### Transcriptomic and proteomic studies

Gene sets of genes with expression correlated with markers of human brain development were obtained from Supplementary Table 12 of the BrainSpan publication^13^. Gene sets of adult brain cell-type markers were obtained from Supplementary Table 3 of Darmanis et al.^10^ and Supplementary Table 5 of Lake et al.^11^, and those for fetal brain cell-type markers were obtained from Supplementary Tables 5 and 6 of Nowakowski et al.^9^ The list of genes with detectable transcription in the adult human brain was obtained from the Human Protein Atlas project^14,15^. Genes expressed in the fetal brain were obtained from GSE104276^16^. The list of proteins that change in the schizophrenia synaptosomal proteome were obtained from Supplementary Table 4 of the original publication^17^.

##### Inferred brain and disease-related gene sets

We included significant genes identified by GWAS studies published by the Psychiatric Genomics Consortium (PGC), for which data are provided on the PGC website (<https://www.med.unc.edu/pgc/results-and-downloads/>). We excluded Alcohol Dependence, Eating Disorders, Obsessive Compulsive Disorder, Tourette's syndrome and Post-Traumatic Stress Disorder (PTSD) as fewer than twenty significant genes were identified in these studies. We obtained lists of significant genes for autism spectrum disorder^18^, major depression disorder^19^, and bipolar disorder^20^ from supplementary tables of the corresponding studies (see Supplementary Table 10 for details). For schizophrenia, we used the list of high-confidence genes published by the PsychENCODE Integrative Analysis^5^ (INT-17 from <http://resource.psychencode.org/>); these are genes supported by three or more evidence sources.

The general list of genes associated with "abnormality of the nervous system" were obtained from the Human Phenotype Ontology^21^. The ontology tree (OBO format) was downloaded from <https://raw.githubusercontent.com/obophenotype/human-phenotype-ontology/master/hp.obo>

on 20^th^ June, 2019 and child terms of the term "HP:0000707 – abnormality of the nervous system" were included.

Genes from all inferred gene sets above were additionally required to have detectable gene expression in the adult or in the fetal brain. For the adult brain, the list of 14,518 "brain-detectable genes" was downloaded from proteinatlas.org^15^ (search results for "tissue_detectable_rna:cerebral cortex;yes"). For the fetal brain, the list of normalized TPM values was downloaded from ftp://ftp.ncbi.nlm.nih.gov/geo/series/GSE104nnn/GSE104276/suppl/GSE104276_all_pfc_2394_UMI_TPM_NOERCC.xls.gz (^16^). Detectable genes are those with TPM greater than zero in at least 20 samples.

#### Supplementary Tables

Supplementary Table 1. Sample and SNP count breakdown by microarray platform

Supplementary Table 2. Regions of high linkage disequilibrium, from which SNPs were excluded

Supplementary Table 3. Description of neurocognitive phenotypes tested and sample breakdown by label

Supplementary Table 4. Correlation of phenotypes with age and sex before and after regression. The Bonferroni-corrected alpha is 0.0056

Supplementary Table 5. Brain-related gene sets

Supplementary Table 6. SNPs associated with binarized PNC neurocognitive phenotypes. SNPs are annotated by functional relevance, tissue and disease expression, eQTL

Supplementary Table 7. Functional annotation for genes associated with neurocognitive variation. Annotated by expression in disease and brain development, and known drug interactions

Supplementary Table 8. Pathways enriched in genetic variants associated with PNC phenotypes (q < 0.05

Supplementary Table 9. Leading edge genes of enriched pathways (q<0.05) and associated SNPs

Supplementary Table 10. Annotations for leading edge genes in pathways (q < 0.05)

Supplementary Table 11. Brain-related gene sets enriched in genetic variants associated with PNC phenotypes (q < 0.05)

Supplementary Table 12. Leading edge genes of enriched brain-related gene sets (q<0.05) and associated SNPs

Supplementary Table 13. Annotations for leading edge genes in brain-related gene sets (q < 0.05)

Supplementary Table 14. Non-coding RNA associated with top SNPs for each task

### Supplementary Figures

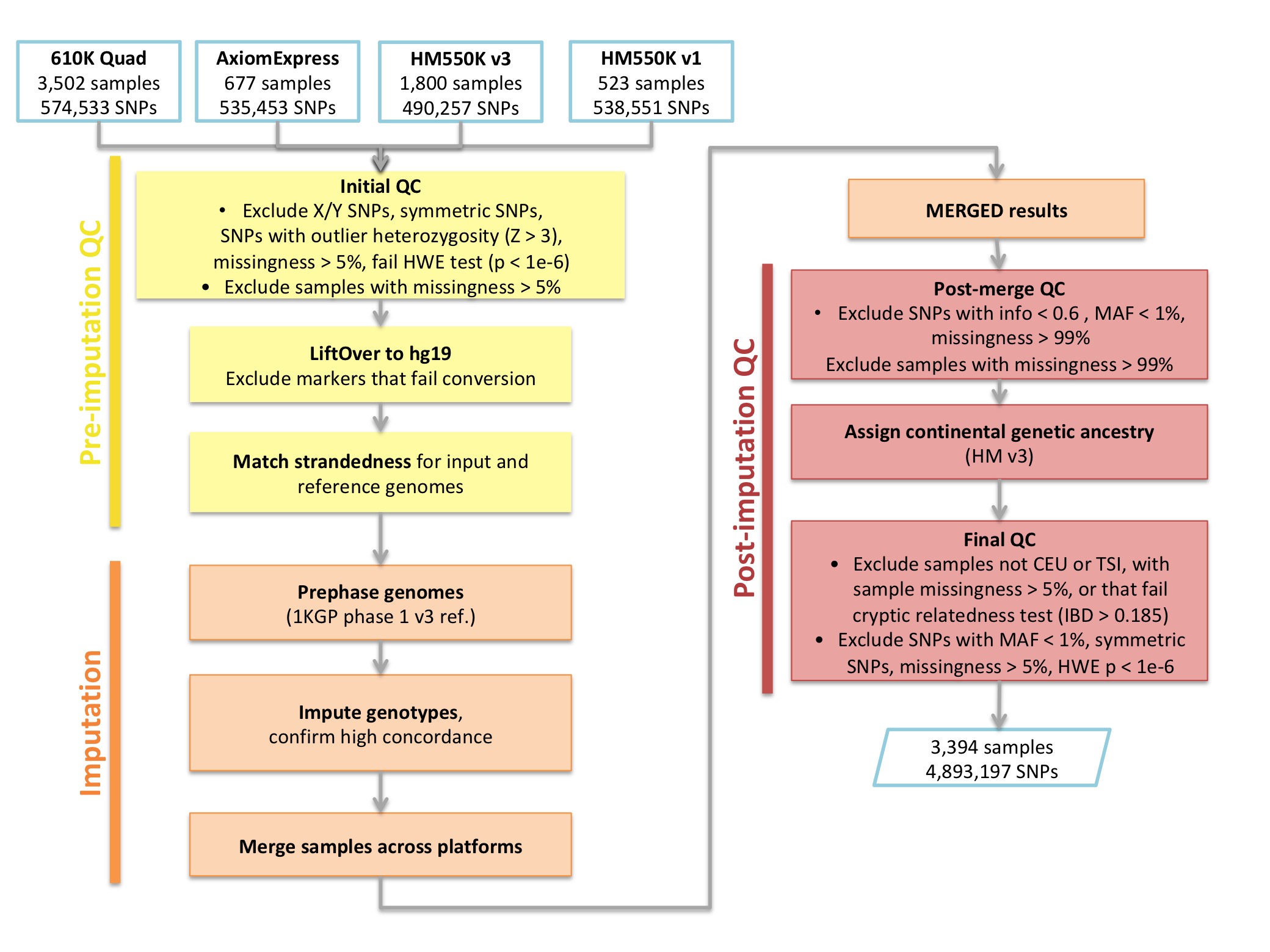

**Supplementary Figure 1.** Workflow for imputation pipeline. Methods after Verma et al. ^22^

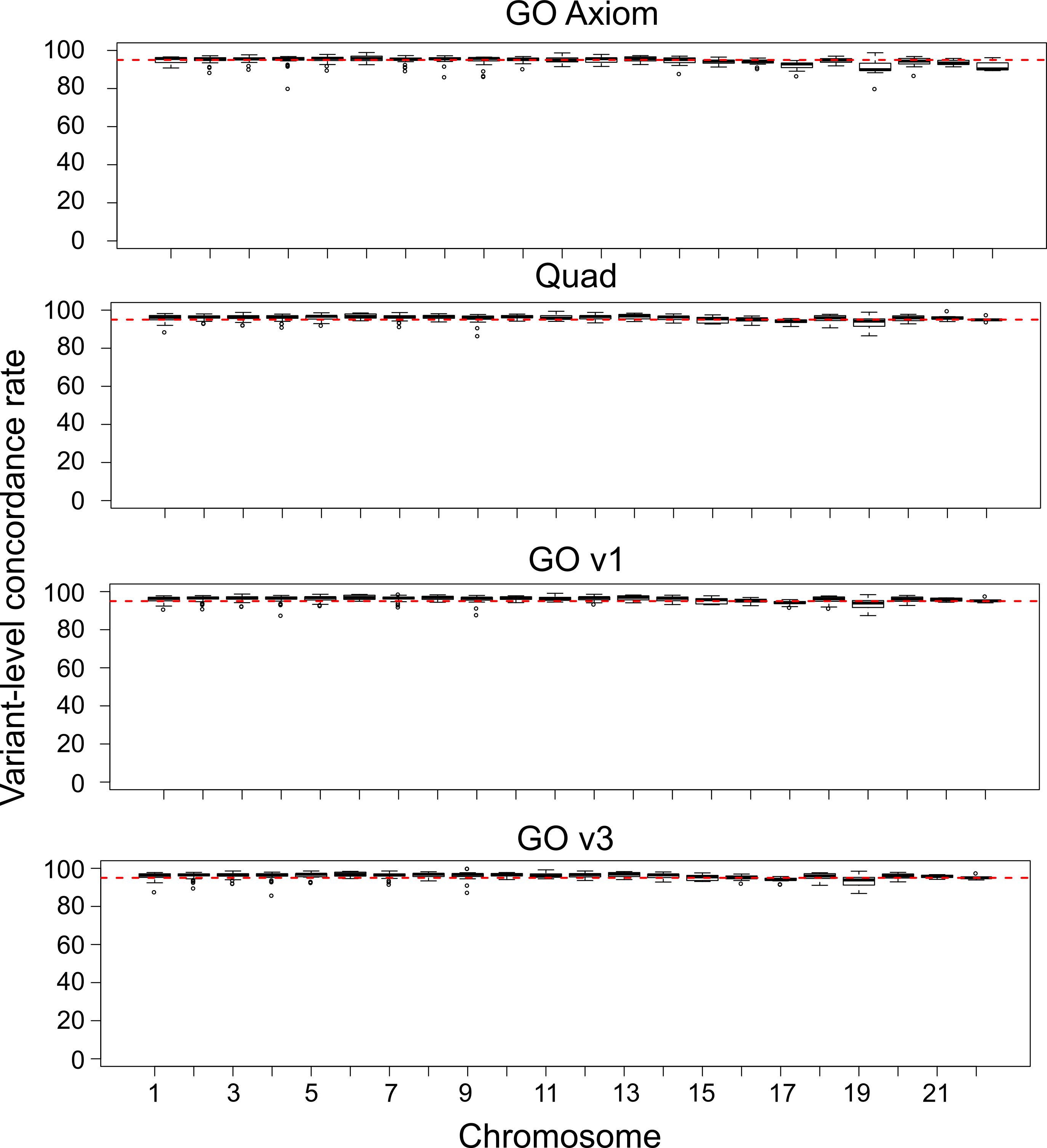

**Supplementary Figure 2.** Concordance rates for imputed SNPs from four microarray platforms comprising of the PNC genotype data. Each panel shows data for one of the microarray platforms, and each boxplot shows distribution of SNP-level concordance levels for a given chromosome (x-axis).

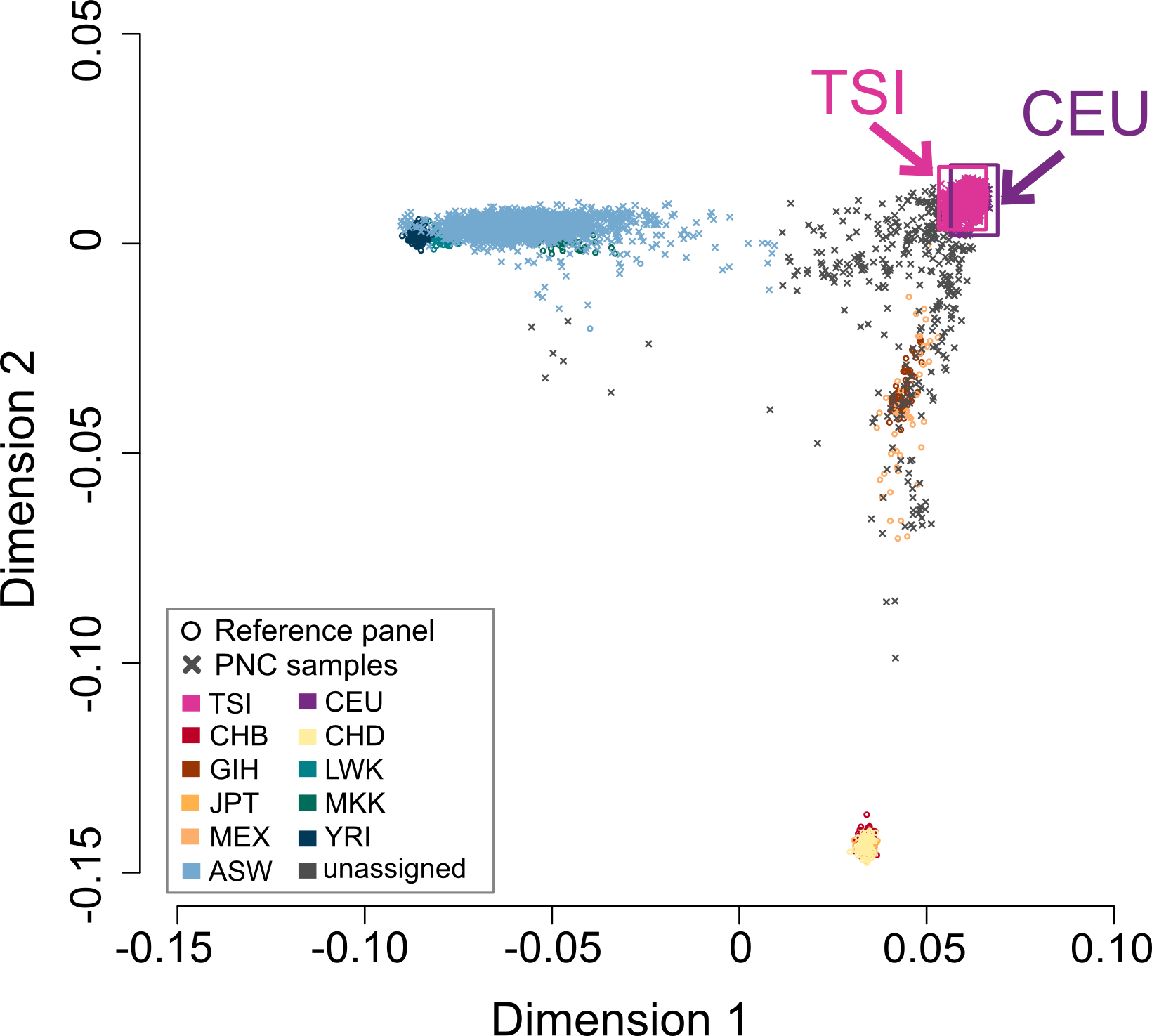

**Supplementary Figure 3.** MDS plot of PNC samples alongside HapMap3 reference panels. Circles show reference samples, and "X" symbols indicate PNC samples. For association analyses, we included individuals with 5SD of the centroid of the CEU and TSI populations (solid box).

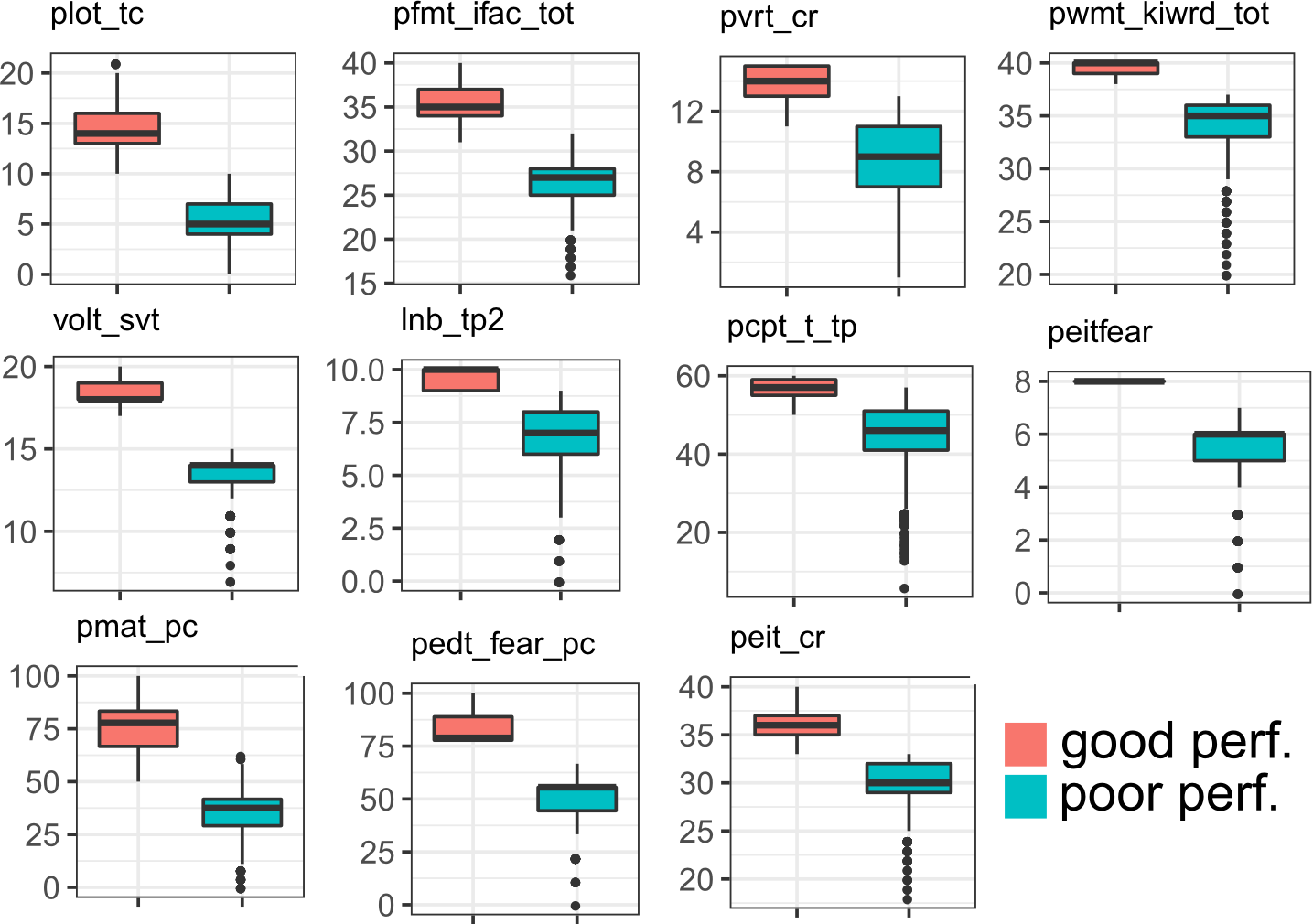

**Supplementary Figure 4.** Untransformed CNB phenotypes, separated by binarized labels

Each panel shows one of the phenotypes tested for pathway enrichment, after the patients were separated into the top and bottom 33% (good vs. poor performance). Units are either “% accuracy” or “number of accurate trials”.

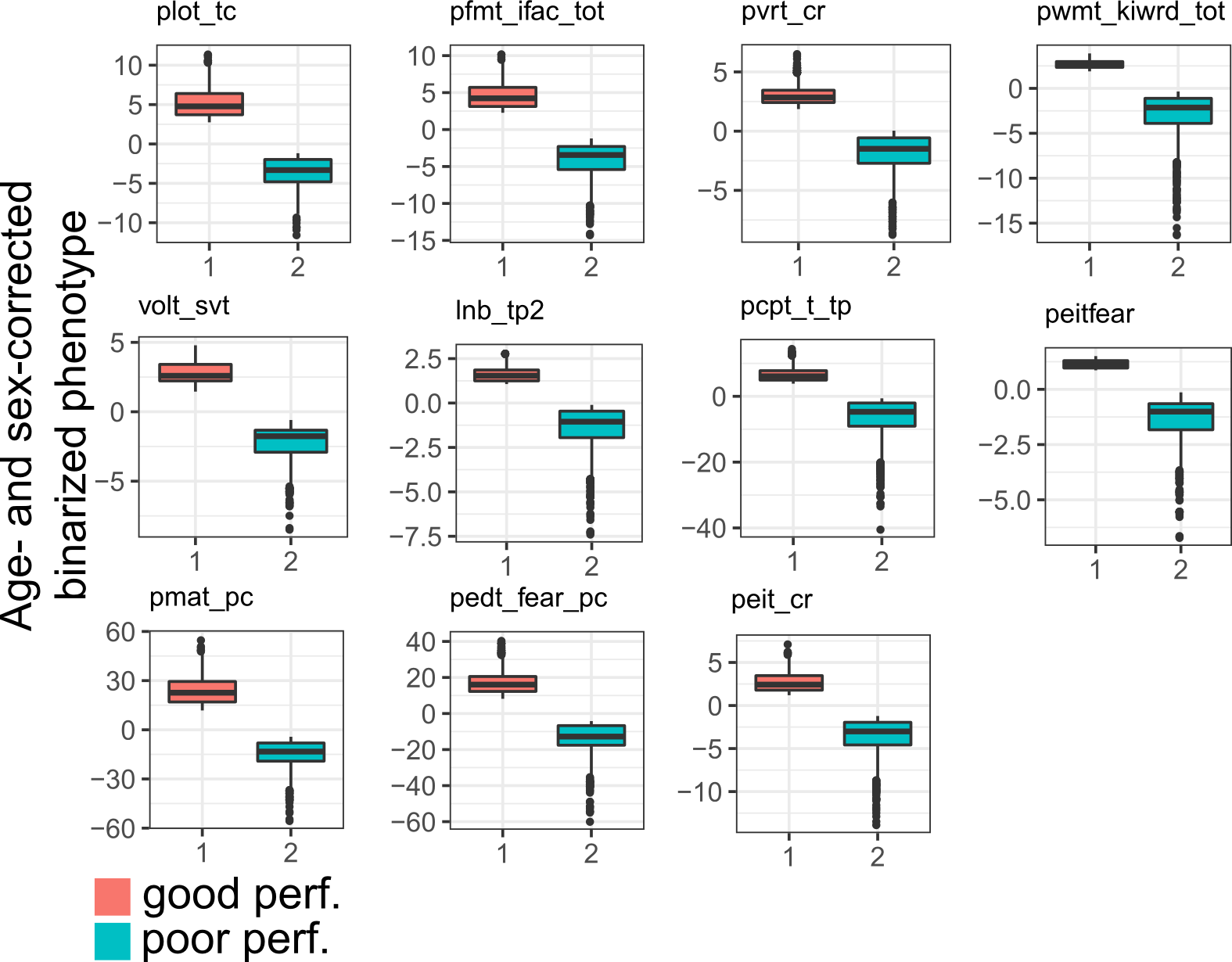

**Supplementary Figure 5.** Phenotype corrected for age and sex, binarized for pathway analysis.

Each panel shows binarized phenotypes after age and sex were regressed out.

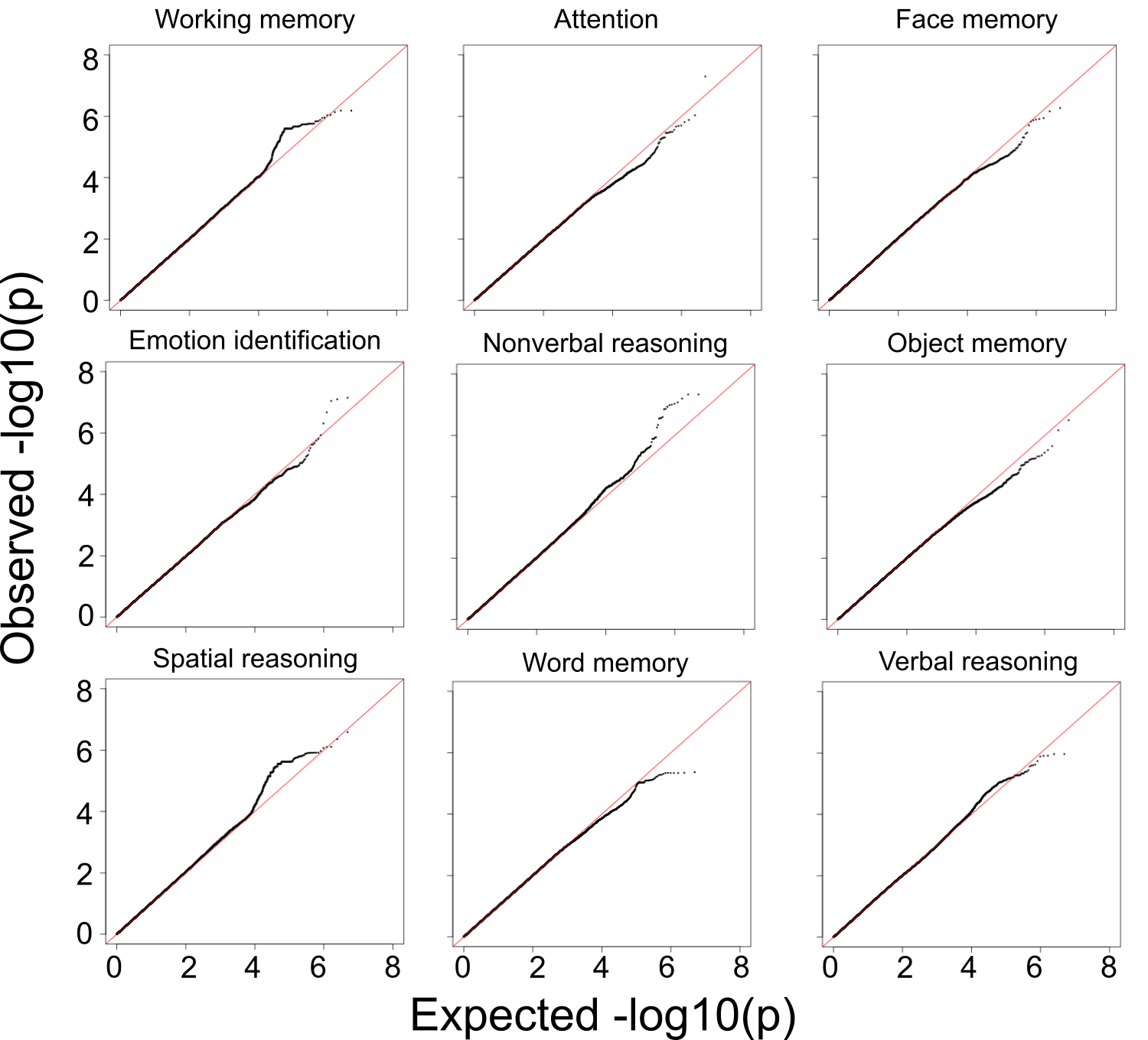

**Supplementary Figure 6.** QQplots for genome-wide association analyses for binarized neurocognitive test performance (GCTA).

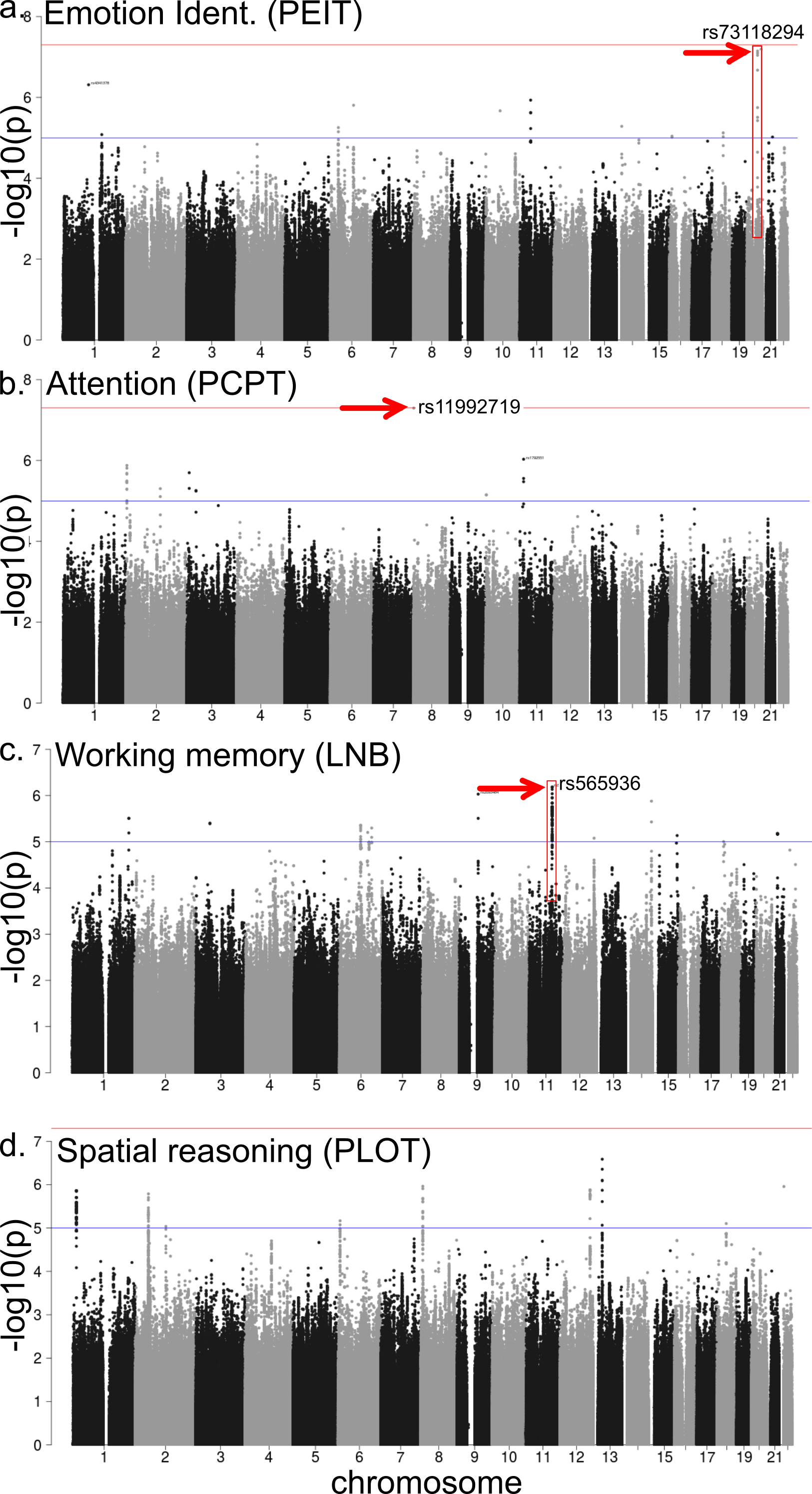

**Supplementary Figure 7.** Manhattan plots for CNB phenotypes with strong or suggestive genotypes. Each panel shows the results for a different phenotype, with the top SNP indicated with an arrow. Where the SNP is part of a larger region of top-ranking SNPs, the region is highlighted with a red rectangle. Note that y-axis scales for the plots differ.

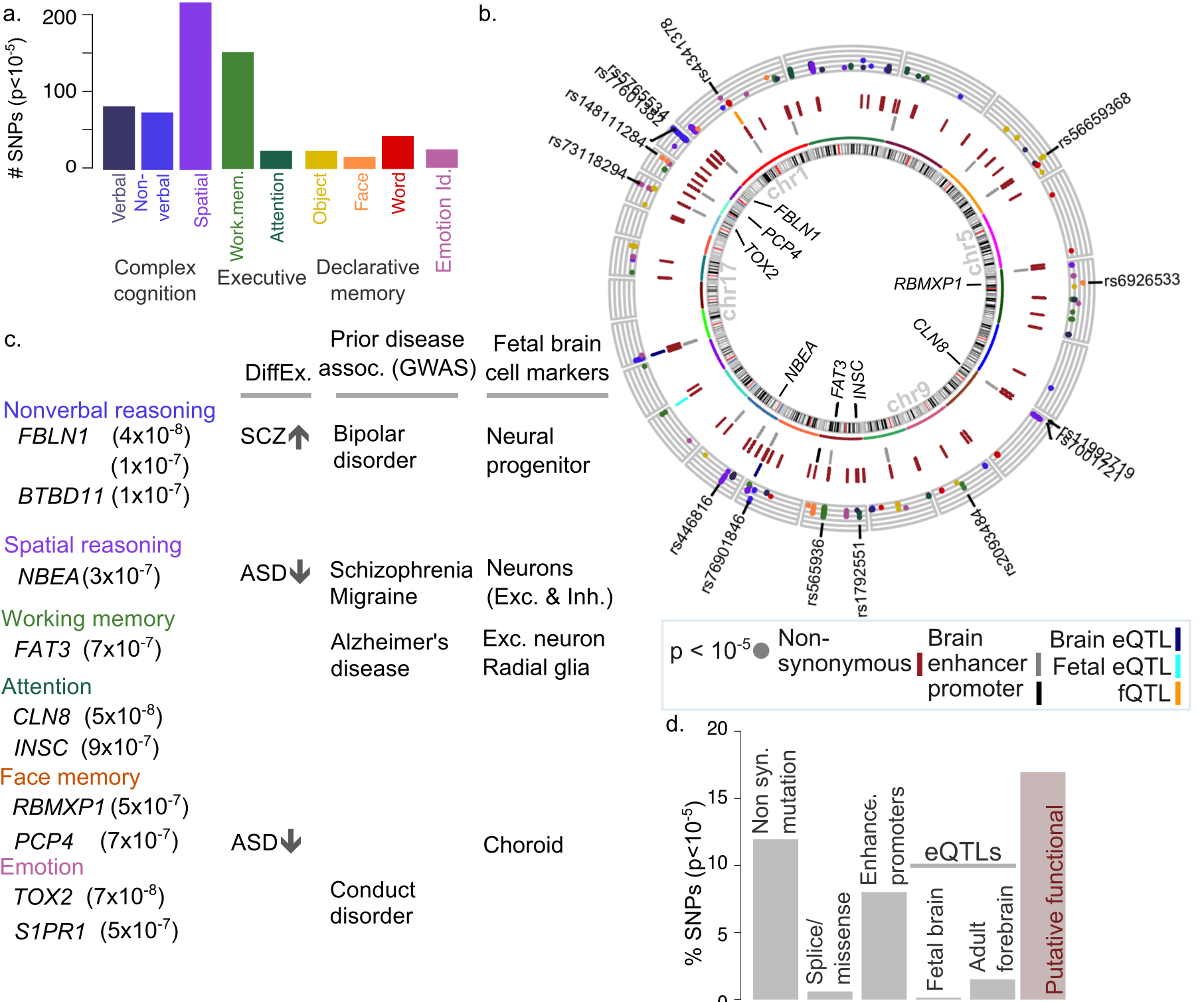

**Supplementary Figure 8.** Genome-wide association analysis for neurocognitive phenotypes from the Philadelphia Neurodevelopmental Cohort.

a. Breakdown of SNPs achieving suggestive significance (p < 10^-5^), by phenotype (top).

b. Suggestive and significant SNPs and associated genes. The outermost ring shows the location of suggestive peaks (p < 10^-5^), coloured by phenotype (see b); y-axis shows –log10(SNP p), so that SNPs with stronger significance are higher. SNPs with p<10^-7^ are labeled. The tracks with ticks indicate functional consequences of associated SNPs. The track closest to the middle indicate SNPs overlapping brain enhancers (light gray) or promoters (black). The dark red middle track indicates SNPs with nonsynonymous variation, including NMD transcript, missense or splice variants (BioMart)^6^. The outermost track indicates QTL associations, including eQTL in adult prefrontal cortex (dark blue), fetal brain (cyan), or neuronal cell proportions in the adult brain (fQTL; orange) (GTEx{Battle, 2017 #26}). Genes associated with top SNPs are indicated within the circle. See Supplementary Note 1 for annotation sources.

c. Genes associated with top SNPs (p < 3x10^-7^) with prior knowledge about relevance to brain development and psychiatric disorders. Columns indicate differential expression in neurodevelopmental disorders^5^ (SCZ = schizophrenia; ASD= autism), significant association with a nervous system disorder^7^, or status as marker gene for specific cell types in fetal brain^9^.

d. Breakdown of functional consequence of top SNPs and by functional consequence (bottom). Consequence shown is limited to effect on protein sequence^6^, presence in enhancers or promoters in adult cortical regions^1^, eQTL in fetal brain, or adult forebrain. Final bar shows cumulative proportion of putatively functional SNPs.

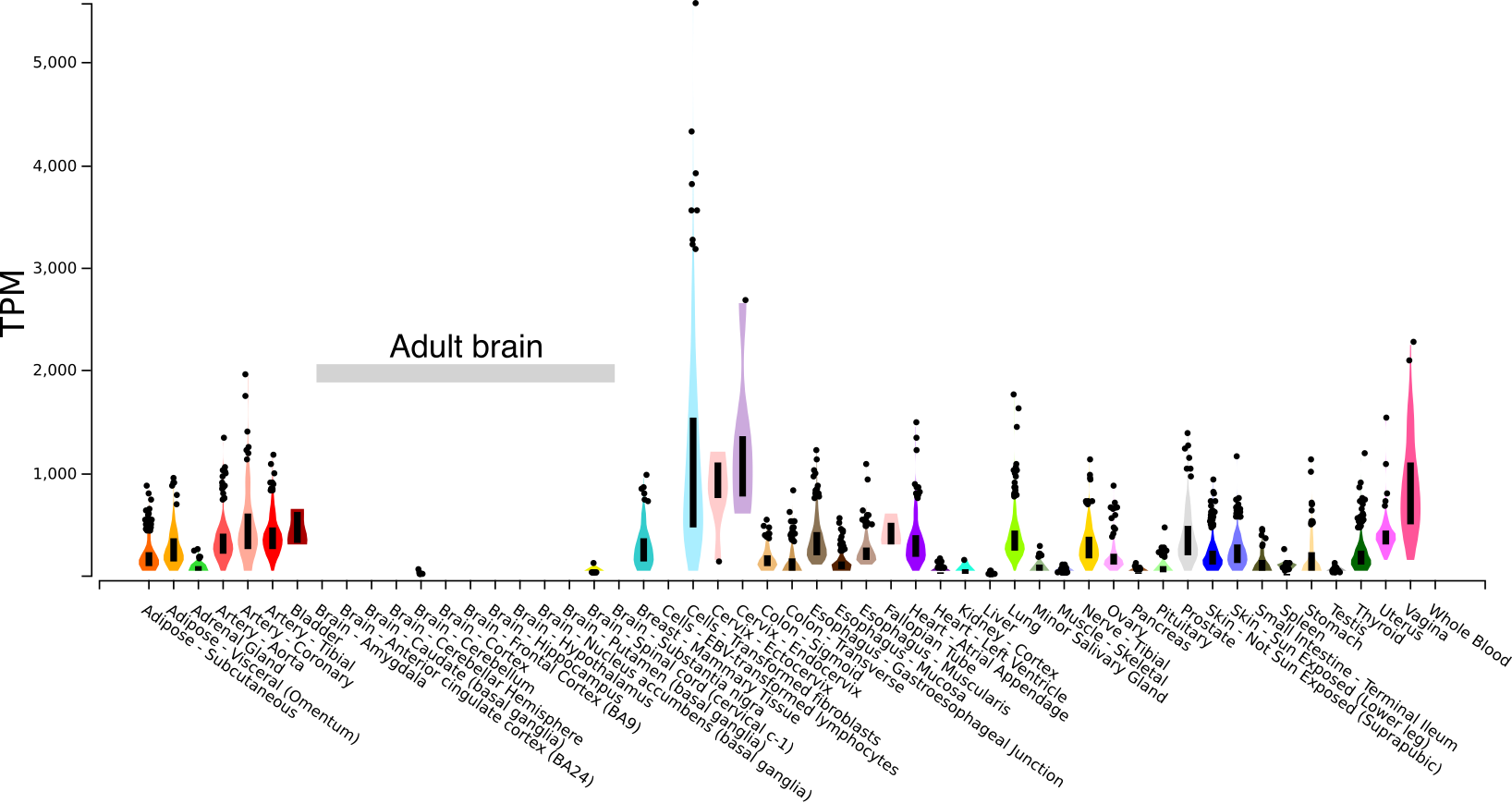

**Supplementary Figure 9.** FBLN1 is expressed at very low levels in the adult human brain, consistent with expression in the fetal brain and subsequent downregulation.

Data and plot from the GTEx portal ([https://www.gtexportal.org](https://www.gtexportal.org/); ^2^).

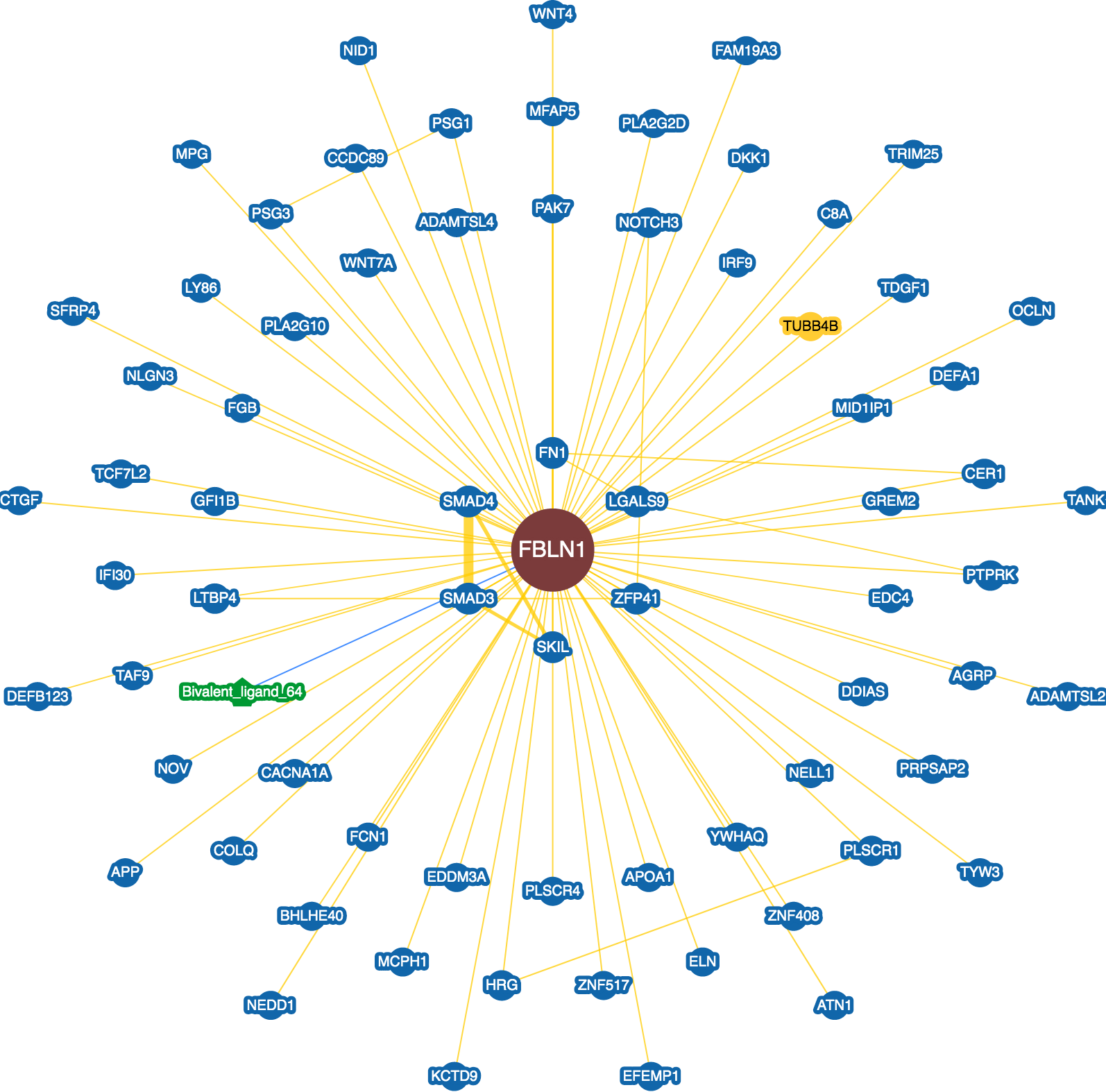

**Supplementary Figure 10.** Physical interactions for FBLN1, from BioGRID^23^.

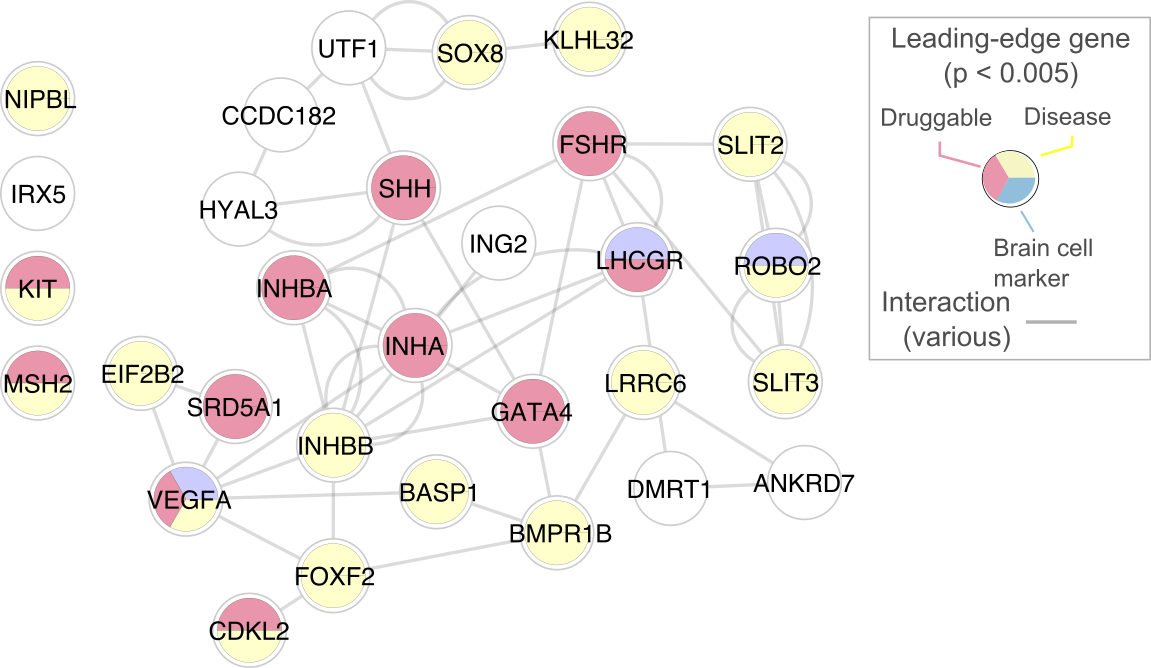

**Supplementary Figure 11.** Leading-edge genes in working memory pathways.

Top leading edge genes in pathways significant in enrichment analysis (pathways with q < 0.05). Only genes with p < 5e-3 are shown (30 genes). Nodes show genes, and fill indicates association with brain cell types, drugs or nervous system disorders (white indicates absence of association). Edges indicate known interactions (GeneMANIA^24^). Full list of genes and attributes in Supplementary Table 10.

### References

1	Kundaje, A. *et al.* Integrative analysis of 111 reference human epigenomes. *Nature* **518**, 317-330, doi:10.1038/nature14248 (2015).

2	Battle, A., Brown, C. D., Engelhardt, B. E. & Montgomery, S. B. Genetic effects on gene expression across human tissues. *Nature* **550**, 204-213, doi:10.1038/nature24277 (2017).

3	Fromer, M. *et al.* Gene expression elucidates functional impact of polygenic risk for schizophrenia. *Nature neuroscience* **19**, 1442-1453, doi:10.1038/nn.4399 (2016).

4	O'Brien, H. E. *et al.* Expression quantitative trait loci in the developing human brain and their enrichment in neuropsychiatric disorders. *Genome biology* **19**, 194, doi:10.1186/s13059-018-1567-1 (2018).

5	Wang, D. *et al.* Comprehensive functional genomic resource and integrative model for the human brain. *Science (New York, N.Y.)* **362**, doi:10.1126/science.aat8464 (2018).

6	Smedley, D. *et al.* The BioMart community portal: an innovative alternative to large, centralized data repositories. *Nucleic acids research* **43**, W589-598, doi:10.1093/nar/gkv350 (2015).

7	Buniello, A. *et al.* The NHGRI-EBI GWAS Catalog of published genome-wide association studies, targeted arrays and summary statistics 2019. *Nucleic acids research* **47**, D1005-d1012, doi:10.1093/nar/gky1120 (2019).

8	Carvalho-Silva, D. *et al.* Open Targets Platform: new developments and updates two years on. *Nucleic acids research* **47**, D1056-d1065, doi:10.1093/nar/gky1133 (2019).

9	Nowakowski, T. J. *et al.* Spatiotemporal gene expression trajectories reveal developmental hierarchies of the human cortex. *Science (New York, N.Y.)* **358**, 1318-1323, doi:10.1126/science.aap8809 (2017).

10	Darmanis, S. *et al.* A survey of human brain transcriptome diversity at the single cell level. *Proceedings of the National Academy of Sciences of the United States of America* **112**, 7285-7290, doi:10.1073/pnas.1507125112 (2015).

11	Lake, B. B. *et al.* Neuronal subtypes and diversity revealed by single-nucleus RNA sequencing of the human brain. *Science (New York, N.Y.)* **352**, 1586-1590, doi:10.1126/science.aaf1204 (2016).

12	Cotto, K. C. *et al.* DGIdb 3.0: a redesign and expansion of the drug-gene interaction database. *Nucleic acids research* **46**, D1068-d1073, doi:10.1093/nar/gkx1143 (2018).

13	Kang, H. J. *et al.* Spatio-temporal transcriptome of the human brain. *Nature* **478**, 483-489, doi:10.1038/nature10523 (2011).

14	Uhlen, M. *et al.* Proteomics. Tissue-based map of the human proteome. *Science (New York, N.Y.)* **347**, 1260419, doi:10.1126/science.1260419 (2015).

15	Yu, N. Y. *et al.* Complementing tissue characterization by integrating transcriptome profiling from the Human Protein Atlas and from the FANTOM5 consortium. *Nucleic acids research* **43**, 6787-6798, doi:10.1093/nar/gkv608 (2015).

16	Zhong, S. *et al.* A single-cell RNA-seq survey of the developmental landscape of the human prefrontal cortex. *Nature* **555**, 524-528, doi:10.1038/nature25980 (2018).

17	Velasquez, E. *et al.* Synaptosomal Proteome of the Orbitofrontal Cortex from Schizophrenia Patients Using Quantitative Label-Free and iTRAQ-Based Shotgun Proteomics. *Journal of proteome research* **16**, 4481-4494, doi:10.1021/acs.jproteome.7b00422 (2017).

18	Grove, J. *et al.* Identification of common genetic risk variants for autism spectrum disorder. *Nature genetics* **51**, 431-444, doi:10.1038/s41588-019-0344-8 (2019).

19	Howard, D. M. *et al.* Genome-wide meta-analysis of depression identifies 102 independent variants and highlights the importance of the prefrontal brain regions. *Nature neuroscience* **22**, 343-352, doi:10.1038/s41593-018-0326-7 (2019).

20	Stahl, E. A. *et al.* Genome-wide association study identifies 30 loci associated with bipolar disorder. *Nature genetics* **51**, 793-803, doi:10.1038/s41588-019-0397-8 (2019).

21	Kohler, S. *et al.* Expansion of the Human Phenotype Ontology (HPO) knowledge base and resources. *Nucleic acids research* **47**, D1018-d1027, doi:10.1093/nar/gky1105 (2019).

22	Verma, S. S. *et al.* Imputation and quality control steps for combining multiple genome-wide datasets. *Frontiers in Genetics* **5**, 370 (2014).

23	Stark, C. *et al.* BioGRID: a general repository for interaction datasets. *Nucleic acids research* **34**, D535-539, doi:10.1093/nar/gkj109 (2006).

24	Franz, M. *et al.* GeneMANIA update 2018. *Nucleic acids research* **46**, W60-w64, doi:10.1093/nar/gky311 (2018).

1	Kundaje, A. *et al.* Integrative analysis of 111 reference human epigenomes. *Nature* **518**, 317-330, doi:10.1038/nature14248 (2015).

2	Battle, A., Brown, C. D., Engelhardt, B. E. & Montgomery, S. B. Genetic effects on gene expression across human tissues. *Nature* **550**, 204-213, doi:10.1038/nature24277 (2017).

3	Fromer, M. *et al.* Gene expression elucidates functional impact of polygenic risk for schizophrenia. *Nature neuroscience* **19**, 1442-1453, doi:10.1038/nn.4399 (2016).

4	O'Brien, H. E. *et al.* Expression quantitative trait loci in the developing human brain and their enrichment in neuropsychiatric disorders. *Genome biology* **19**, 194, doi:10.1186/s13059-018-1567-1 (2018).

5	Wang, D. *et al.* Comprehensive functional genomic resource and integrative model for the human brain. *Science (New York, N.Y.)* **362**, doi:10.1126/science.aat8464 (2018).

6	Smedley, D. *et al.* The BioMart community portal: an innovative alternative to large, centralized data repositories. *Nucleic acids research* **43**, W589-598, doi:10.1093/nar/gkv350 (2015).

7	Buniello, A. *et al.* The NHGRI-EBI GWAS Catalog of published genome-wide association studies, targeted arrays and summary statistics 2019. *Nucleic acids research* **47**, D1005-d1012, doi:10.1093/nar/gky1120 (2019).

8	Carvalho-Silva, D. *et al.* Open Targets Platform: new developments and updates two years on. *Nucleic acids research* **47**, D1056-d1065, doi:10.1093/nar/gky1133 (2019).

9	Nowakowski, T. J. *et al.* Spatiotemporal gene expression trajectories reveal developmental hierarchies of the human cortex. *Science (New York, N.Y.)* **358**, 1318-1323, doi:10.1126/science.aap8809 (2017).

10	Darmanis, S. *et al.* A survey of human brain transcriptome diversity at the single cell level. *Proceedings of the National Academy of Sciences of the United States of America* **112**, 7285-7290, doi:10.1073/pnas.1507125112 (2015).

11	Lake, B. B. *et al.* Neuronal subtypes and diversity revealed by single-nucleus RNA sequencing of the human brain. *Science (New York, N.Y.)* **352**, 1586-1590, doi:10.1126/science.aaf1204 (2016).

12	Cotto, K. C. *et al.* DGIdb 3.0: a redesign and expansion of the drug-gene interaction database. *Nucleic acids research* **46**, D1068-d1073, doi:10.1093/nar/gkx1143 (2018).

13	Kang, H. J. *et al.* Spatio-temporal transcriptome of the human brain. *Nature* **478**, 483-489, doi:10.1038/nature10523 (2011).

14	Uhlen, M. *et al.* Proteomics. Tissue-based map of the human proteome. *Science (New York, N.Y.)* **347**, 1260419, doi:10.1126/science.1260419 (2015).

15	Yu, N. Y. *et al.* Complementing tissue characterization by integrating transcriptome profiling from the Human Protein Atlas and from the FANTOM5 consortium. *Nucleic acids research* **43**, 6787-6798, doi:10.1093/nar/gkv608 (2015).

16	Zhong, S. *et al.* A single-cell RNA-seq survey of the developmental landscape of the human prefrontal cortex. *Nature* **555**, 524-528, doi:10.1038/nature25980 (2018).

17	Velasquez, E. *et al.* Synaptosomal Proteome of the Orbitofrontal Cortex from Schizophrenia Patients Using Quantitative Label-Free and iTRAQ-Based Shotgun Proteomics. *Journal of proteome research* **16**, 4481-4494, doi:10.1021/acs.jproteome.7b00422 (2017).

18	Grove, J. *et al.* Identification of common genetic risk variants for autism spectrum disorder. *Nature genetics* **51**, 431-444, doi:10.1038/s41588-019-0344-8 (2019).

19	Howard, D. M. *et al.* Genome-wide meta-analysis of depression identifies 102 independent variants and highlights the importance of the prefrontal brain regions. *Nature neuroscience* **22**, 343-352, doi:10.1038/s41593-018-0326-7 (2019).

20	Stahl, E. A. *et al.* Genome-wide association study identifies 30 loci associated with bipolar disorder. *Nature genetics* **51**, 793-803, doi:10.1038/s41588-019-0397-8 (2019).

21	Kohler, S. *et al.* Expansion of the Human Phenotype Ontology (HPO) knowledge base and resources. *Nucleic acids research* **47**, D1018-d1027, doi:10.1093/nar/gky1105 (2019).

22	Verma, S. S. *et al.* Imputation and quality control steps for combining multiple genome-wide datasets. *Frontiers in Genetics* **5**, 370 (2014).

23	Stark, C. *et al.* BioGRID: a general repository for interaction datasets. *Nucleic acids research* **34**, D535-539, doi:10.1093/nar/gkj109 (2006).

24	Franz, M. *et al.* GeneMANIA update 2018. *Nucleic acids research* **46**, W60-w64, doi:10.1093/nar/gky311 (2018).
